# Long-read RNA-seq resolves isoform-level and context-specific regulatory architecture of complex traits in cattle

**DOI:** 10.64898/2026.06.04.730050

**Authors:** Weijie Zheng, Qi Zhang, Jinfeng He, Xiaoning Zhu, Mian Gong, Houcheng Li, Lin Liu, Jianbin Li, Zhu Ma, Bo Han, Yu Zhang, Jingsheng Lu, Bingjin Lin, Bingxing An, Zijiao Guo, Yanan Liu, Aixia Du, Di Zhu, Xinfeng Liu, Jianguo Li, Xuemei Lu, Lingzhao Fang, Dongxiao Sun

## Abstract

Traditional short-read RNA-seq lacks the resolution to fully capture full-length isoforms and complex alternative splicing, leaving a "missing regulation" gap in existing molecular atlases. Here, we present a population-scale, multi-omics dissection of bovine complex traits using matched deep whole-genome sequencing (30×), long-read (ONT) and short-read RNA-seq, as well as metabolome data from 432 dairy cows across four distinct lactation stages. We expanded the bovine transcript atlas with 13,177 novel isoforms and fine-mapped 1,088,900 regulatory effects across 11 diverse molecular phenotypes. Notably, long-read mapping uncovered 606 genes with isoform expression quantitative trait loci (12.8%, eQTL) and 1,116 genes with detailed splicing QTL (sQTL) (31.8%) that were missed by conventional short-read mapping. We further showed that lactation-specific regulation was profoundly mediated by the metabolic microenvironment (46.8%) and cellular composition (4.3%). Across genome-wide associations (GWAS) of 90 bovine complex traits, isoform-level and context-specific regulatory effects captured GWAS loci missed by short-read eQTL and sQTL analyses, increasing colocalized loci from 35.2% to 60.3%. These regulatory variants also exhibited evolutionary constraints with human immune and metabolic loci. Our study establishes an invaluable resource (https://cattleblr.farmgtex.org) demonstrating that long-read transcriptomics is essential for resolving the complexity of regulatory mechanisms underlying mammalian physiology.

## Introduction

The central goal of modern genetics is to decode the molecular mechanisms by which genomic variation translates into complex phenotypes. While genome-wide association studies (GWAS) have identified thousands of variants associated with complex traits in both humans and livestock^1–3^, approximately 90% reside in noncoding regions, hindering their functional interpretation^4^. Despite large-scale efforts like Genotype-Tissue Expression Project (GTEx) and Farm animal (FarmGTEx) have mapped thousands of steady average gene expression quantitative trait loci (eQTL) in humans and livestock species^5–11^, respectively, a significant "missing regulation" gap persists, where the majority of trait-associated variants cannot be reconciled with steady-state gene-level regulatory signals^12,13^.

This disconnect is partially attributable to the technical limitations of conventional short-read RNA sequencing (srRNA-seq). By shearing transcripts into small fragments, srRNA-seq aggregates fragmented reads to estimate gene-level expression, while inferring local alternative splicing based on junction reads clustering^14,15^. This inevitably collapses transcript diversity and obscures full-length isoform structures. Long-read RNA-seq (lrRNA-seq) overcomes these constraints by capturing full-length transcripts^15^, enabling accurate isoform-level quantification and high-resolution characterization of complex alternative splicing architectures^16–18^. Recent studies in humans have demonstrated that transcript-level regulation frequently operates independently of steady-state gene expression and may explain previously unresolved GWAS loci^18,19^. However, population-scale studies using lrRNA-seq remain limited, particularly in farmed animals, with small sample sizes restricting analyses largely to allele-specific analysis rather than population-based quantitative trait loci (QTL) mapping^20^.

Furthermore, current molecular atlases largely treat genetic regulation as static process, overlooking evidence that genetic effects are highly context dependent and can be reshaped by developmental, metabolic, immune and endocrine environments^21–23^. In particular, many trait-associated variants exhibit context-specific regulatory activity that cannot be captured under steady-state conditions^4,24^. Lactation represents a good biological system for investigating dynamic genetic regulation, as it involves profound and coordinated remodeling of metabolism, immunity and endocrine signaling. We therefore hypothesized that a substantial fraction of context-dependent genetic regulation operates through isoform-level mechanisms that remain unresolved by conventional gene-expression analyses.

To address these limitations, we conducted a population-scale, multi-omics dissection of the bovine functional transcriptome as part of the FarmGTEx initiative^9^, by generating matched deep whole-genome sequencing (mean 35.6×), srRNA-seq, lrRNA-seq by Oxford Nanopore sequencing (ONT), and untargeted metabolome profiles from 432 individuals across four distinct lactation stages, including dry (n = 112), early (n = 124), peak (n = 117), and late (n = 79) stages (**Fig. 1a-c; Supplementary Table 1**). Leveraging both srRNA-seq and lrRNA-seq, we substantially enhanced the bovine transcript atlas and defined 11 types of molecular phenotypes, including two expression-based phenotypes (i.e., gene expression, isoform expression), and nine splicing-based phenotypes (i.e., splicing, isoform usage ratio, and seven types of alternative splicing events, such as skipped exon, SE; alternative 5′ splice site, A5′; alternative 3′ splice site, A3′; retained intron, RI; mutually exclusive exons, MX; alternative first exon, AS; and alternative last exon, AL) **(Fig. 1c).** We systematically mapped regulatory variants of these molecular phenotypes and explored their lactation-specific regulatory patterns. Furthermore, we assessed how these regulatory effects were mediated by metabolic and cellular microenvironments **(Fig. 1 d, e)**. By integrating GWAS results of 90 complex cattle traits and 52 complex human traits, selection signatures between dairy and beef cattle, and cross-species comparative analyses, we revealed the temporal and evolutionary dimensions of gene regulation underlying complex and adaptive traits in cattle and humans **(Fig. 1f)**. In summary, our work establishes the valuable long-read CattleGTEx resource with lactation stage resolution (https://cattleblr.farmgtex.org) for understanding the multi-layered genetic control of mammalian physiology, offering insights for both precision agriculture and evolutionary medicine.

**Fig. 1.**
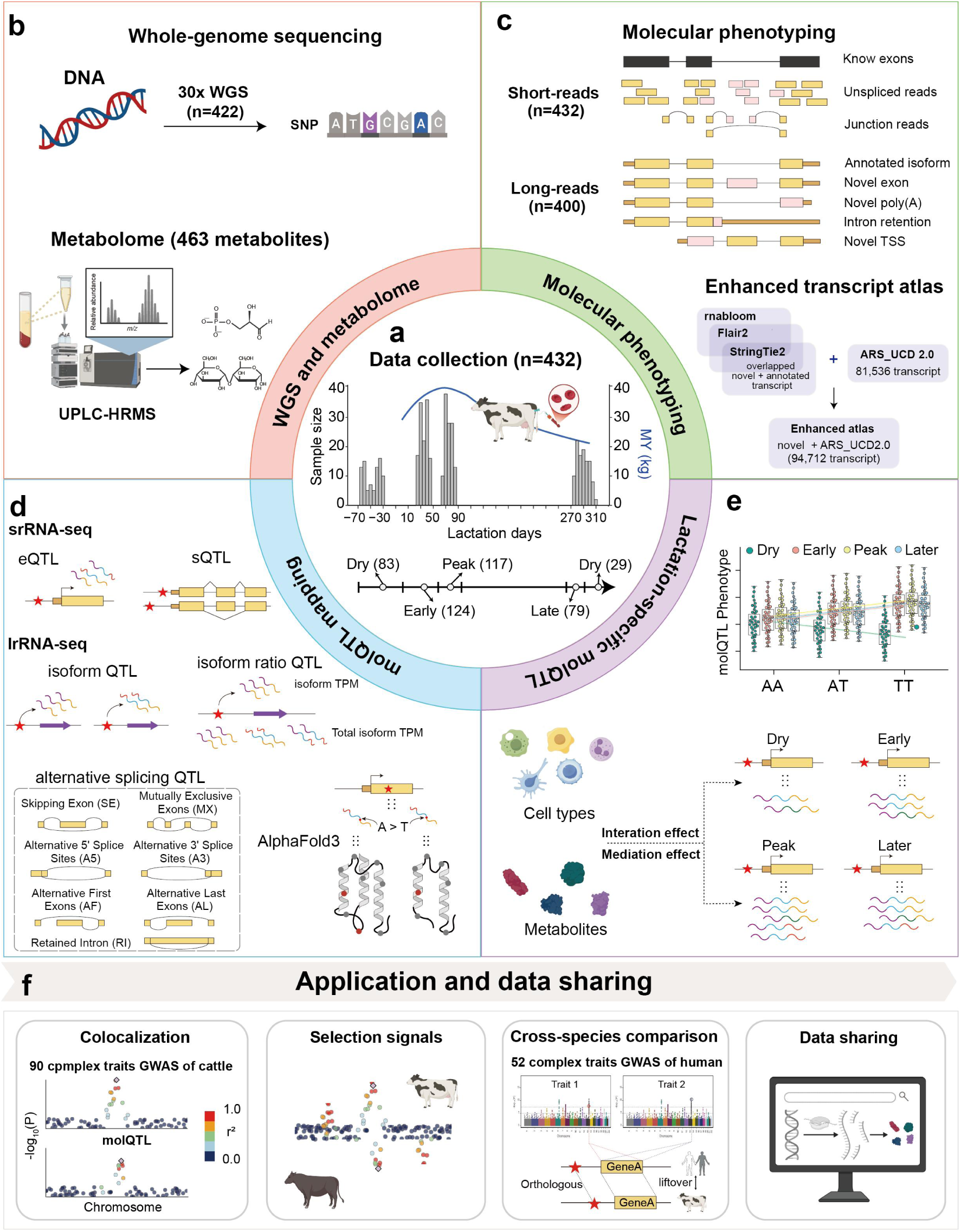
Experimental design and analytical framework of the long-read blood CattleGTEx resource. **a**, Peripheral blood were collected from 432 dairy cattle across four lactation stages, including 112 dry, 124 early-lactation, 117 peak-lactation and 79 late-lactation individuals. **b**, Whole-genome sequencing (30×) and serum untargeted metabolomic profiling were performed on the same animals. **c**, Matched short-read and long-read RNA-seq were generated from peripheral blood. Transcriptomes were reconstructed via integrative assembly using RNA-Bloom, FLAIR2 and StringTie2, and were merged with the ARS-UCD2.0 to generate an enhanced transcriptome atlas. **d**, Using this resource, cis-expression QTL (eQTL) and splicing QTL (sQTL) were mapped from short-read RNA-seq data, whereas isoform expression QTL (isoQTL), isoform ratio QTL (irQTL) and seven classes of alternative splicing QTL (asQTL) were identified from long-read RNA-seq. The predicted effects of variants on protein structure were further evaluated using AlphaFold3. **e**, Lactation stage-specific molQTL were identified, and the mediating effects of blood cell composition and metabolite abundance on stage-specific regulation were systematically investigated. **f**, The resource was applied in GWAS of dairy complex traits, population selection scans between dairy and beef cattle, and cross-species evolutionary analyses between cattle and humans. All datasets are publicly available.

## Results

### Long-read RNA-seq reveals extensive isoform diversity in bovine blood transcriptome

In total, we obtained 34.35 and 4.68 billion clean reads with a mean uniquely mapping rate of 96.3% and 85.0% for srRNA-seq and lrRNA-seq, respectively **(Supplementary Table 2-3)**. Among all lrRNA-seq reads, 83.8% represented full-length transcripts, with a mean read length of 1,080bp compared to 143 bp for short reads, providing the structural resolution necessary to define complex splicing events (**Extended Data Fig. 1 a-b**).

Using a hybrid transcript assembly approach (**see Methods**), we identified 29,284 candidate isoforms that were supported by at least two of three methods, including Stringtie2^25^, Flair2^26^ and RNAbloom^27^ **(Fig. 2a; Extended Data Fig. 1c)**. After stringent quality control with SQANTI3^28^, we retained 25,196 high-confidence isoforms across 11,151 genes **(Supplementary Table 4)**. Strikingly, 13,177 of these isoforms derived from 6,018 genes were absent from the current ARS_UCD2.0 reference genome annotation **(Extended Data Fig. 1c; Supplementary Fig. 1)**. Among the novel isoforms, 56.7% were classified novel isoform with at least a new splicing site (novel not in catalog, NNC), and 27.9% as novel isoform with a new combination of known splice sites (novel in catalog, NIC), indicating that lrRNA-seq substantially expands the known transcriptomic landscape compared to previous short-read efforts **(Extended Data Fig. 1d,e)**. Coding potential analysis using four independent tools (i.e., TransDecoder^29^, CPC2^30^, CNCI^31^, and CPAT^32^) further categorized these novel isoforms into 9,523 protein-coding and 3,654 non-coding transcripts **(Extended Data Fig. 1f,g)**. Alternative splicing analysis revealed that alternative 3′ splice sites (24.5%), exon skipping (23.5%), and alternative 5′ splice sites (21.6%) were the dominant drivers of this diversity, whereas alternative last exons were rare (1.5%; **Fig. 2b; Supplementary Table 5)**. To generate an enhanced bovine blood transcript atlas, we finally merged the 13,177 novel isoforms with the 81,536 transcripts from the ARS-UCD2.0 annotation, resulting in 94,712 isoforms from 38,387 genes **(Extended Data Fig. 1h,i)**.

**Fig. 2.**
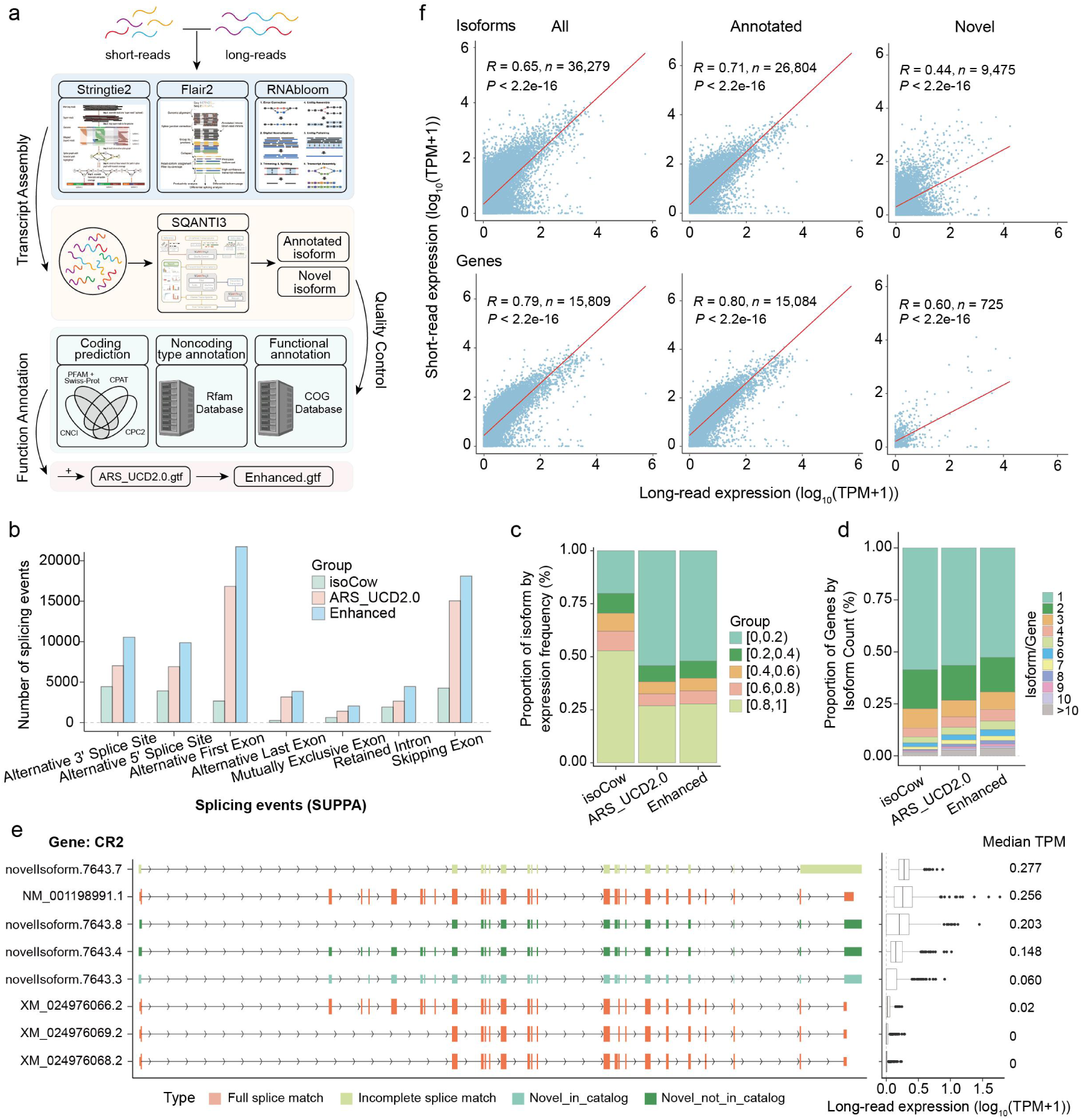
Hybrid transcriptome reconstruction integrating long- and short-read RNA sequencing. **a**, Overview of the analytical framework: (Ⅰ) hybrid transcriptome assembly with RNA-Bloom, FLAIR2 and StringTie2; (Ⅱ) isoform quality control and classification with SQANTI3; (Ⅲ) coding potential of novel isoforms assessed using CNCI, CPAT, CPC2 and TransDecoder; functional annotation via the COG database. **b**, Numbers of alternative splicing events identified by SUPPA in the reference (ARS_UCD2.0), isoCow (novel Isoforms) and enhanced annotations. **c**, Fraction of individuals (n = 400) expressing each isoform (TPM > 0.1), grouped into five bins with 0.2 intervals, where higher proportions (>0.2) indicate more ubiquitously expressed. **d**, Distribution of isoforms count per gene in the three transcriptome panels. **e**, Isoform diversity at the *CR2* locus. Left panel is a schematic diagram of the structure of the novel isoform and ARS_UCD2.0 isoforms. Right panel shows the log_10_(TPM+1) of each isoform measured by long-read RNA-seq across 400 individuals. Box plots indicate the interquartile range (IQR) with the median, and whiskers extend to 1.5× IQR. **f**, Pearson correlation of median TPM quantified by long- and short-read RNA-seq for all, annotated, and novel isoforms and genes.

Furthermore, we quantified expression levels of all isoforms in the enhanced atlas across the 400 cows, and revealed that 79.8% (10,510) of the novel isoforms were robustly expressed (>20% of individuals, Transcripts Per Million, TPM > 0.1), a significantly higher proportion than the 45.8% observed for the original ARS-UCD2.0 isoforms **(Fig. 2c)**. In addition, this expanded catalog increased the proportion of genes expressing multiple isoforms from 43.5% to 47.3% **(Fig. 2d)**. For instance, while the ARS-UCD2.0 annotation contains four isoforms of *CR2*, only one of which reached our expression threshold, long-read sequencing detected four additional novel isoforms, three of which were highly abundant (median TPM > 0.1; **Fig. 2e**). *CR2* is a critical mediator of B-cell activation^33^. These findings demonstrate that lrRNA-seq identifies functionally relevant transcripts that are missed by current annotations, setting the stage for a higher-resolution dissection of the regulatory variants governing these molecular phenotypes. In addition, median expression levels for both genes and isoforms showed high correlation between short- and long-read data **(Fig. 2f)**, supporting the use of lrRNA-seq for accurate transcript- and gene-level quantification^34^. Leveraging this enhanced atlas, we quantified 11 molecular phenotypes across the 400 cows, including 17,176 gene expression and 86,780 splicing events identified from srRNA-seq, as well as nine more refined isoform-level features derived from lrRNA-seq, including 33,451 isoform expression, 33,451 isoform usage ratios, 7,481 SE, 6,435 AF, 5,247 A3′, 4,701 A5′, 2,061 RI, 1,414 AL, and 855 ME **(Supplementary Fig. 2)**.

### Long-read isoQTL mapping captures extensive regulatory variation beyond gene-level eQTL

To identify genetic variants regulating the diverse molecular phenotypes detected above, we first characterized genomic variants of the 422-cow cohort using WGS, identifying 9.94 million common SNPs (minor allele frequency, MAF > 0.05) **(Extended Data Fig. 2a-e; Supplementary Table 6)**. Most variants were located in intronic (66.73%) and intergenic (19.46%) regions, with only 0.11% affecting splice-sites (**Extended Data Fig. 2f)**. These variants broadly covered across 10 previously predicted chromatin states (58.1-96.5%)^35^, with ∼7,000 variants on average within ±1 Mb of transcription start sites (TSS) used for subsequent *cis*-QTL mapping (**Extended Data Fig. 2 g-k**).

After accounting for potential confounding factors, we performed *cis*-molQTL mapping using Omiga^36^ for all 11 molecular phenotypes, identifying 13,836 molGenes (79.72% of all tested genes) with at least one significant variant **(Extended Data Fig. 3; Supplementary Fig. 3-4).** For expression-based QTL, we identified 12,330 eGenes for gene expression (71.79%) and 4,751 isoGenes for isoform expression (55.76%). Notably, 606 (12.76%) isoGenes were not eGenes **(Fig. 3a)**. Of these, 63.7% showed significance only at the single-isoform level; 15.2% reflected opposing effects across different isoforms, with 47.92% of isoform were low expression (isoform ratio < 25%); 14.4% harbored dispersed, independent signals across isoforms; and 2.8% were diluted by highly expressed isoforms lacking effects **(Fig. 3b-f)**. Fine-mapping further identified 469,954 and 359,738 candidate SNPs in eGenes and isoGenes, respectively, of which 74.76% and 67.02% were novel **(Supplementary Fig. 5)**. Functional annotation showed that novel isoVariants were preferentially enriched at splice_acceptor_variant and splice_donor_variant sites, indicating that isoform-level mapping captured regulatory variants primarily involved in splice-site usage and isoform selection **(Fig. 3g)**. In contrast, novel eVariants were more enriched in coding-related annotations, including initiator_codon_variant, start_lost, stop_gained and stop_lost, as well as in bivEnh and ReprPC chromatin states, suggesting potential effects on protein-coding capacity, transcript stability and regulatory chromatin contexts **(Fig. 3g-h)**.

**Fig. 3.**
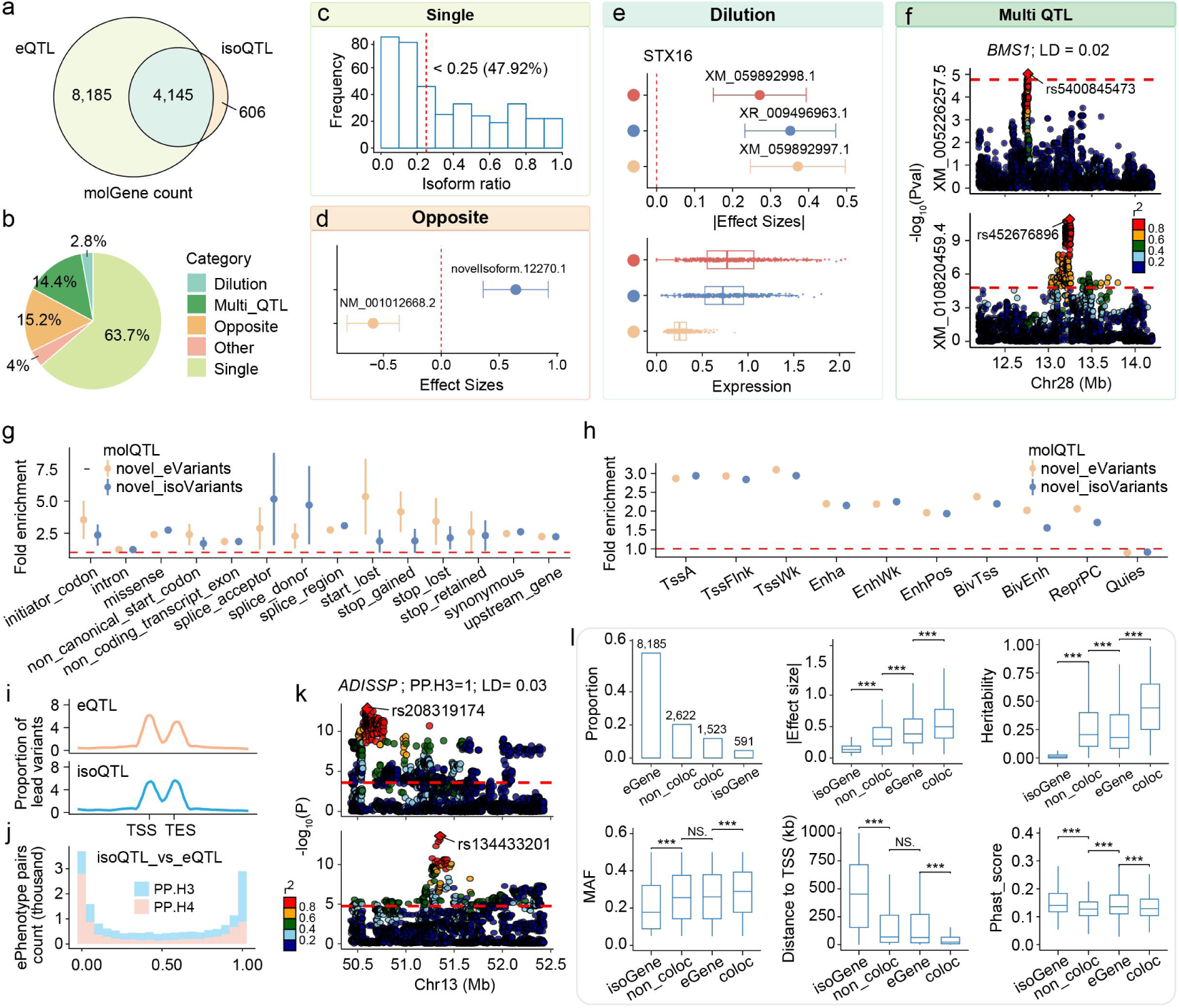
Long-read isoQTL mapping reveals transcript-level regulatory heterogeneity beyond gene-level eQTL. **a,** Overlap between eGenes and isoGenes. **b**, Proportion of mechanistic categories underlying the identification of novel isoGenes. **c-f,** Representative regulatory patterns explaining why transcript-level molQTL may be missed or attenuated at the gene level. **(c)** Distribution of expression ratios of the QTL-associated isoform among isoGenes in which only one isoform showed a significant QTL signal. These transcript-level regulatory effects may be diluted at the gene level by other highly expressed transcripts without significant QTL signals. **(d)** Opposite regulatory effects across transcripts from the same gene result in signal cancellation. **(e)** Regulatory-effect strength is inversely related to transcript abundance, leading to dilution at the gene level. **(f)** Multiple molecular phenotypes from the same gene are regulated by independent signals, resulting in mutually diluted gene-level effects. **g-h,** Enrichment of novel fine-mapped variants across 14 sequence ontology categories **(g)** and 10 chromatin states **(h)**. **i**, Distribution of lead variants of molGenes relative to TSS and TES. **j,** Co-localization results between isoPhenotypes and ePhenotypes. PP.H3 indicates molecular phenotypes driven by two independent causal variants, whereas PP.H4 indicates shared causal variant. **k,** Representative example of molecular phenotypes regulated by independent cis-regulatory loci. **l,** Comparison of eGene-specific, isoGene-specific, colocalized and non-colocalized signals in terms of their proportion, absolute regulatory effect size (|β|), *cis-h²*, MAF, distance to the TSS and PhastCons conservation score. *P* values were calculated using two-sided paired Wilcoxon signed-rank tests. NS: non-significant; *: *P* < 0.05; **: *P* < 0.01; ***: *P* < 0.001.

To further explore regulatory relationships across molecular layers, we observed that both eQTL and isoQTL were concentrated near transcription start site (TSS) **(Fig. 3i)**. However, only 27% of isoQTL colocalized with eQTL for the same gene (PP.H4 > 0.8), suggesting extensive specific regulation across regulatory layers **(Fig. 3j; Supplementary Table 7)**. A representative example is *ADISSP*, in which transcript-level regulation was driven by lead variants independent of those regulating overall gene expression **(Fig. 3k)**. In addition, genes with colocalized signals showed larger effect sizes, higher *cis-h*^2^, expression, and MAF, but lower evolutionary conservation and shorter distances to the TSS compared with genes regulated by a single molphenotype **(Fig. 3l).** These findings demonstrate that long-read isoform-level molQTL mapping captures regulatory variants that are largely missed by conventional gene-level eQTL analyses, providing a more comprehensive view of genetic regulation across transcriptomic layers.

### Long-read splicing QTL mapping improves regulatory resolution of alternative splicing

Conventional short-read RNA-seq infers local alternative splicing mainly from clustered junction reads, resulting in relatively coarse and fuzzy splicing phenotypes. Leveraging the ability of long-read lrRNA-seq to directly quantify isoform usage and specific splicing events, we identified 6,816 sGenes for alternative splicing (56.8%), 3,300 irGenes for isoform usage ratio (38.7%), and multiple classes of event-resolved asGenes, including 328 RI_asGenes (27.4%), 568 AF_asGenes (30.3%), 777 A5_asGenes (27.3%), 1,064 SE_asGenes (27.0%), 269 AL_asGenes (41.4%), 839 A3_asGenes (26.5%) and 107 MX_asGenes (23.6%) **(Extended Data Fig. 3)**. Importantly, many long-read-derived splicing signals were not detected as conventional sGenes, including 1,044 irGenes (31.6%), 240 SE_asGenes (22.6%), 236 A3_asGenes (28.1%), 219 A5_asGenes (28.2%), 154 AF_asGenes (27.1%), 114 RI_asGenes (34.8%), 51 AL_asGenes (19.0%) and 19 MX_asGenes (17.8%) **(Fig. 4a)**. To explore the reasons for the omission of novel asGenes in conventional sQTL mapping, we found that 7.4% showed opposing allelic effects across splicing phenotypes within the same gene, leading to signal cancellation; 21.1% reflected independently regulated splicing phenotypes, and 38.2% were significant only for a single splicing phenotype, resulting in diluted aggregate signals **(Fig. 4b-d)**. Fine-mapping further identified 378,455 sVariants, 241,103 irVariants and 3,552-32,938 event-specific asVariants, of which 64.8%, 58.3% and 44.2-57.7%, respectively, were novel **(Fig. 4e)**. These novel variants were enriched in splice-related annotations, supporting their direct involvement in transcript processing, and were also enriched in active promoter and enhancer states, indicating that transcript-structure variation is shaped by both splice-proximal and canonical cis-regulatory elements **(Fig. 4f-g)**.

**Fig. 4.**
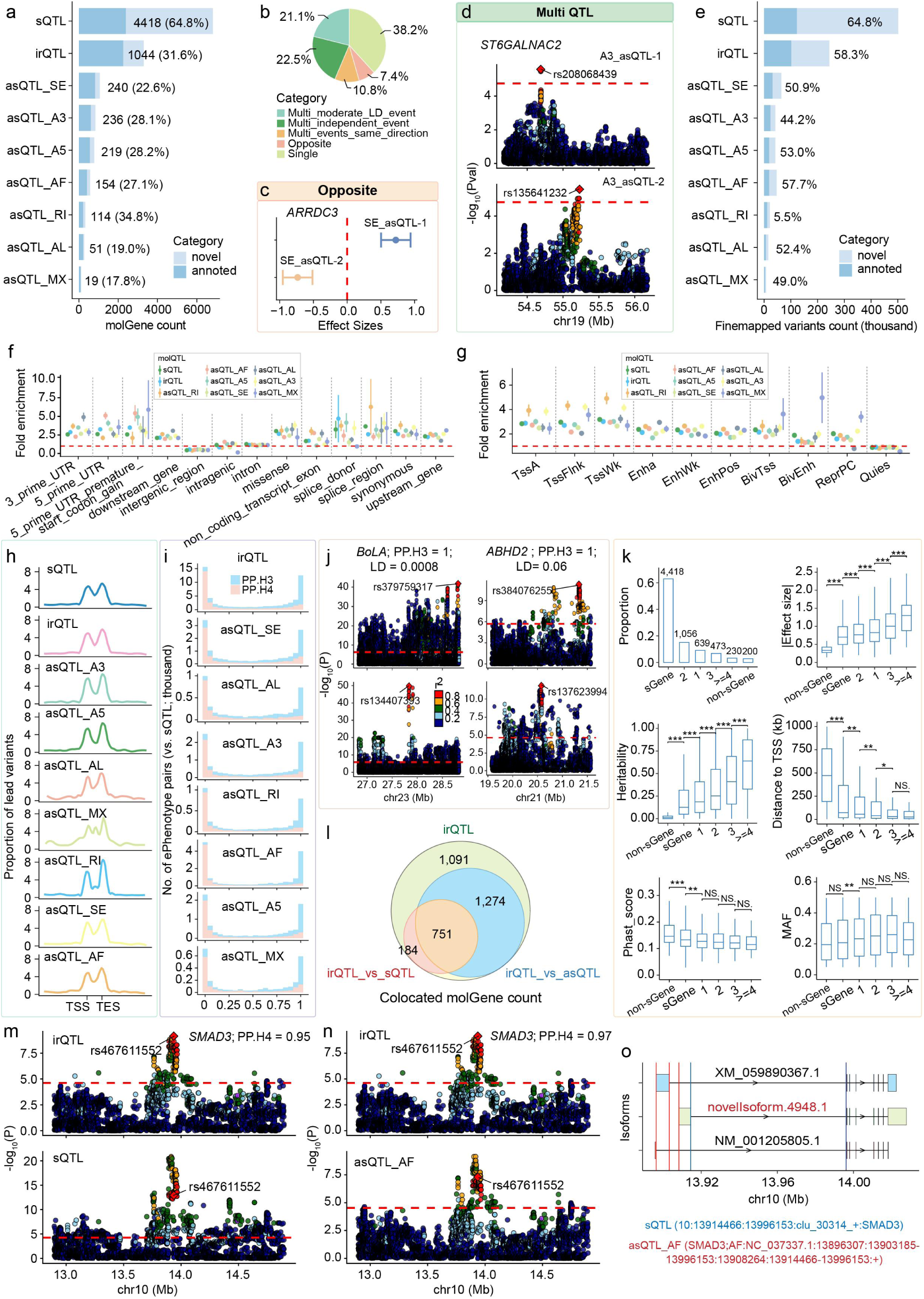
Long-read splicing-based molQTL mapping improves the resolution of splicing regulation. **a,** Number of molGenes identified by splicing-based molQTL mapping. Values indicate the number and proportion of novel molGenes. Novel sGenes were defined as sGenes not detected as asGenes or irGenes, whereas novel asGenes and irGenes were defined as those not detected as conventional sGenes. **b,** Proportion of mechanistic categories underlying the identification of novel asGenes. **c-f,** Representative regulatory patterns explaining why asQTL may be missed or attenuated at the sQTL. **(c),** Opposite regulatory effects across splicing phenotypes from the same gene lead to signal cancellation at the aggregate sQTL level. **(d),** Multiple splicing phenotypes from the same gene are regulated by independent variants, resulting in diluted aggregate sQTL signals. **e,** Number of fine-mapped variants identified across nine splicing-based molecular phenotypes. Values indicate the number and proportion of novel variants, defined according to the same criteria as in **a**. **f-g,** Enrichment of novel fine-mapped variants across 14 sequence ontology categories **(f)** and 10 chromatin states **(g)**. **h**, Distribution of lead variants of molGenes relative to TSS and TES. **i,** Co-localization results between irPhenotypes or asPhenotypes and sPhenotypes. **j,** Representative example of molecular phenotypes regulated by independent cis-regulatory loci. **k,** Comparison of sGene-specific, other molGene-specific, colocalized signals in terms of their proportion, |β|, *cis-h²*, MAF, distance to the TSS and PhastCons conservation score. *P* values were calculated using two-sided paired Wilcoxon signed-rank tests. NS: non-significant; *: *P* < 0.05; **: *P* < 0.01; ***: *P* < 0.001. **l,** Overlap among irGenes whose irQTL colocalized with conventional sQTL and those whose irQTL colocalized with asQTL. **m-o,** *SMAD3* as a representative example in which the irQTL of a specific transcript colocalized with both a conventional sQTL and an AF_asQTL. The AF event specifically gives rise to this isoform, refining the aggregate sQTL signal to a defined splicing event and isoform.

Contrary to expression-based molQTL, splicing-based molQTL were preferentially distributed near transcription end site (TES), particularly for retained intron **(Fig. 4h)**. Colocalization analysis revealed limited sharing with conventional sQTL: only 18.8% of irQTL and 10.7-22.4% of event-specific asQTL colocalized with sQTL (PP.H4 > 0.8) **(Fig. 4i)**. Examples such as *BoLA* and *ABHD2* showed isoform-level regulatory signals driven by variants distinct from those regulating aggregate splicing phenotypes **(Fig. 4j)**. Consistently, colocalized signals showed larger effect sizes, higher *cis-h*^2^, higher expression levels and higher MAF, but lower evolutionary conservation and shorter distances to the TSS compared with signals detected in a splicing phenotype **(Fig. 4k)**.

Finally, we evaluated whether event-resolved asQTL improve interpretation of irPhenotypes regulation. Among 3,300 irGenes, 61.4% colocalized with asQTL, whereas only 28.3% colocalized with conventional sQTL **(Fig. 4l; Supplementary Fig. 6)**, demonstrating that long-read asQTL provide substantially finer mechanistic resolution for isoform-level regulation. A representative example is *SMAD3*, the isoform (novelIsoform.4948.1) colocalized with both a sQTL and an AF_asQTL **(Fig. 4m,n)**. The AF event specifically generated this novel isoform **(Fig. 4o)**, refining an aggregate sQTL association to a defined splicing event and improving mechanistic interpretability.

### Transcription and metabolic reprogramming across lactation stages

While the previous sections defined the global regulatory architecture of the bovine transcriptome, gene regulation is not a static process but one that undergoes profound reprogramming in response to physiological demand. To characterize these dynamics across the lactation cycle **(Fig. 5a)**, we first performed time-series analyses of gene expression. We observed a marked transition from immune activation and muscle morphogenesis during the "dry" (non-lactating) period to lipid response and circulatory development in early and peak lactation. Late lactation was characterized by the upregulation of genes enriched in transporter activity and energy metabolism, reflecting the physiological shift toward sustained nutrient export **(Fig. 5b; Supplementary Fig. 7)**. Weighted gene co-expression network analysis (WGCNA)^37^ further grouped these programs into 19 functional modules, including immune-related modules associated with the dry period, progesterone-related modules in early lactation, and a peak-associated module (M10) enriched in JAK–STAT, mTOR and prolactin signaling pathways, which are key reivers of milk synthesis^38–40^ **(Extended Data Fig. 4)**.

**Fig. 5.**
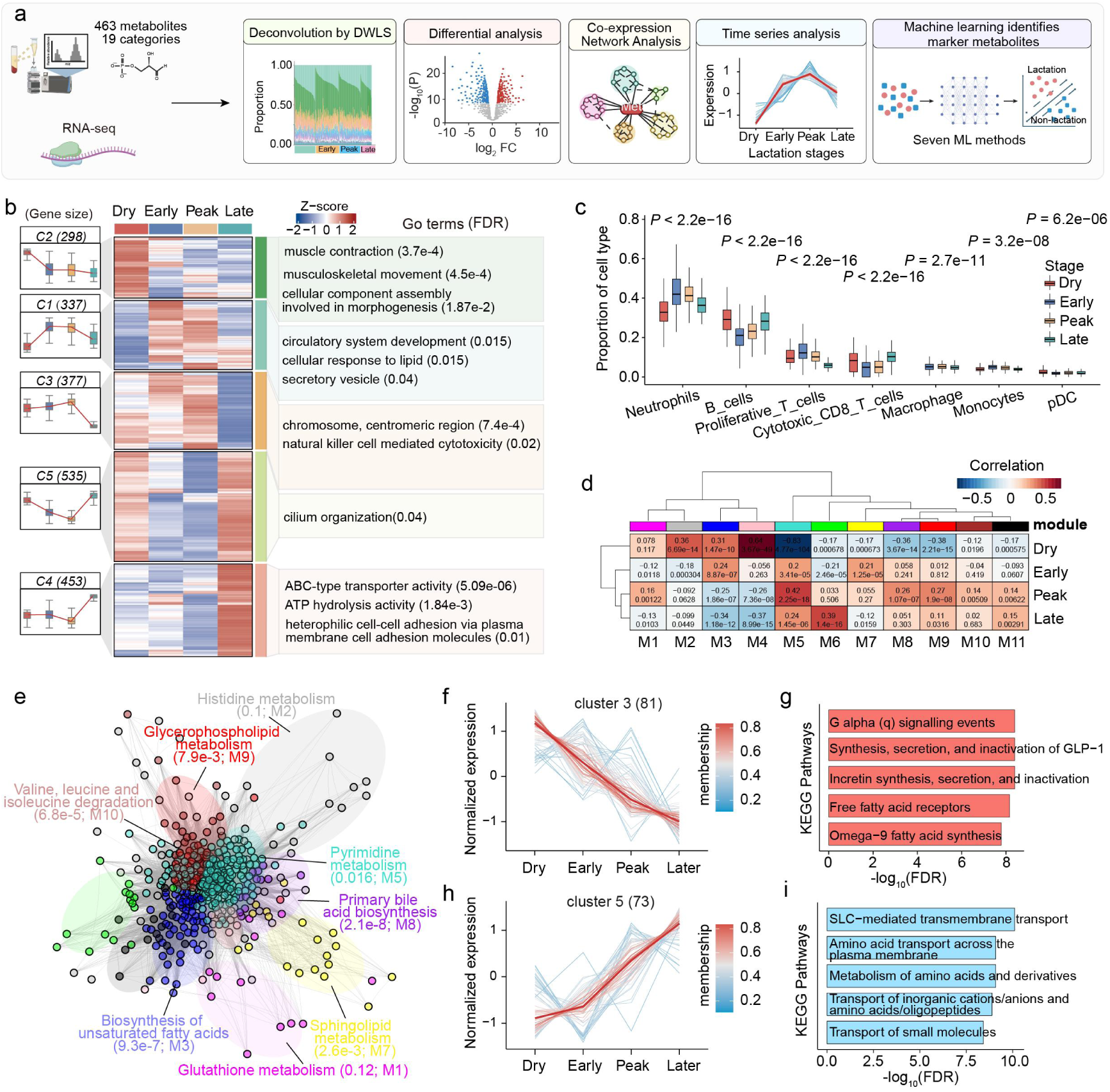
Transcriptional and metabolic reprogramming across lactation stages. **a,** Analytical framework for characterizing lactation-associated transcriptional and metabolic reprogramming. **b**, Time-series analysis identified five distinct expression trends across the four lactation stages. The heatmap shows module-level expression patterns, with representative enriched GO terms and corresponding FDR values shown on the right. **c.** Boxplots showing the proportions of the seven major cell types (mean proportion >2%) across lactation stages. *P* values were calculated using one-way ANOVA. **d**, Co-expression network analysis identified 11 distinct metabolite modules significantly associated with four lactation stages. **e**, KEGG pathway enrichment for the 11 co-expression modules, with each point representing a metabolite, colored by module membership, and lines indicating correlations between metabolites within the same module. **f-i**, KEGG pathway enrichment analysis of the 11 metabolite co-expression modules. Each point represents a metabolite and is colored according to module membership **(f, h)**; lines indicate correlations between metabolites within the same module **(g, i)**.

To explore how changes in the blood cellular composition across lactation stages, we deconvoluted our bulk RNA-seq data using a blood single-cell data comprising 13 major cell types^41^ **(Supplementary Fig. 8a–b).** Our analysis revealed that seven cell types underwent significant proportional shifts across the lactation cycle, with neutrophils and B cells representing the dominant populations. Notably, neutrophils reached highest proportion in early lactation (43.3%) and declined to the lowest level later in lactation (33.3%), whereas B cells showed the inverse trend, rising from 20.4% to 29.9%. This reciprocal pattern indicates a fundamental transition from innate to adaptive immune responses as lactation progresses **(Fig. 5c)**.

Beyond cellular dynamics, lactation was accompanied by extensive metabolic reprogramming. To characterize these dynamics across the lactation cycle, we profiled 463 metabolites across 19 classes in the serum metabolome, of which 416 passed quality control and exhibited clear stage-dependent separation (**Extended Data Fig. 5a,b; Supplementary Table 8**). Among them, 132 metabolites had significant heritability estimates (*P* < 0.05), with an average estimate of 0.32, suggesting moderate-to-high genetic control of serum metabolite variation (**Extended Data Fig. 5c**). Co-expression and time-series analyses revealed coordinated metabolic trajectories, characterized by a decline in lipid- and hormone-related pathways and a progressive increase in amino acid transport and transmembrane transport processes as lactation progressed (**Fig. 5d-i; Extended Data Fig. 5d)**. Through differential metabolite analysis and machine-learning models incorporating SHAP interpretation^42^, we identified the top 20 predictive metabolites that enable accurate discrimination between lactation and non-lactation, including *Thymol Sulfate* and *Proline Betaine* **(Extended Data Fig. 5e)**. Integration with the transcriptomic atlas further revealed a coherent transcriptional-metabolic axis, where genes upregulated in dry-period (e.g., *CXCL12*) were negatively correlated with lactation-associated metabolites (e.g., *Thymol Sulfate*) **(Supplementary Fig.9; Extended Data Fig. 5f)**.

### Lactation-specific regulatory effects

We next sought to determine whether these shifting transcriptional programs are regulated by stage-specific regulatory variants. By performing molQTL mapping for each stage independently, we identified that 5.97–11.30% of molQTL exhibited stage-specific effects **(Fig. 6a)**, including 2,832 lactation-specific eGenes (25.44% of all test genes), 1,095 isoGenes (36.08%), 1,460 sGenes (30.74%), 751 irGenes (34.50%), and 15-177 (24.19%-31.68%) various asGenes. Notably, 937 (85.57%) lactation-specific isoGenes were not lactation-specific eGenes, as well as 642 (85.49%) irGenes and 14-152 (83.33%-94.12%) various asGenes were not lactation-specific sGenes **(Fig. 6b)**, indicating that gene-level analyses substantially underestimated regulatory complexity. For example, the MX_asQTL of *PPARD*, involved in fatty acid metabolism^43,44^, was specifically regulated by rs383139695 at early lactation, while the AF_asQTL of *SMAD3*, the signaling mediator, was regulated by rs110714686 at peak lactation **(Fig. 6c-d)**. Beyond these "on-off" binary effects, we identified variants that were shared across stages but exhibited significant differences in effect size, such as an sQTL (rs135255180) for the immune regulator ***IRF5***, which shows substantially stronger regulatory activity during late lactation compared to peak stage **(Fig. 6e)**.

**Fig. 6.**
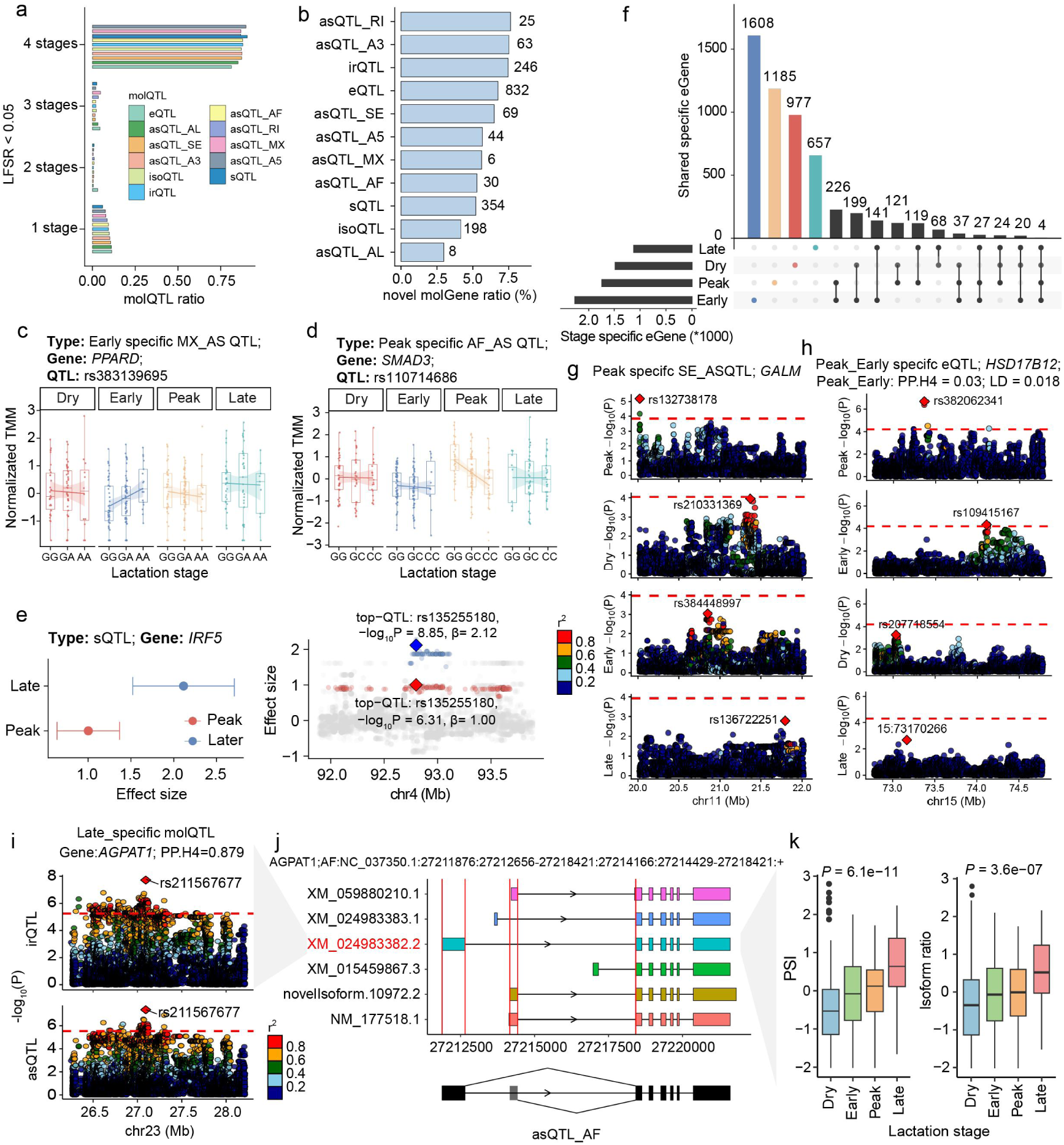
Lactation stage-specific molQTL reveal dynamic regulatory programs. **a**, Proportions of cis-molQTL that are stage-specific or shared across two, three or all four lactation stages. **b,** Number and proportion of novel molGenes regulated by lactation stage-specific molQTL, compared with those identified by standard molQTL mapping. **c,d**, Two representative examples of lactation stage-specific molQTL. **e**, Example of molQTL shared between two stages with distinct effect sizes, as indicated by non-overlapping confidence intervals. **f**, UpSet plot showing the overlap of all stage-specific molGenes. **g-h**, There are two modes of stage-specific regulation: regulated only by a single lactation stage-specific signal **(g)**; controlled by independent signals from multiple lactation stages **(h)**. **i**, Examples of stage-specific regulation: isoform usage ratio of XM_024983382.2 and alternative first exons event (AF) phenotype of *AGPAT1* regulated by the same SNP at late lactation. **j**, AF event generates the specific isoform. **k**, Box plots comparing the PSI of AF event (left) and the usage ratio of isoform (right) among four lactation stage. *P* values were assessed using one-way ANOVA.

By integrating signals across multiple molecular layers, we discovered that 31.52% of all genes were dynamically regulated across the lactation cycle. The majority of these dynamic regulations (81.78%) were confined to a single stage, such as the peak-lactation-specific splicing control of *GALM*, a gene essential for hexose metabolism **(Fig. 6f,g)**. The remaining genes were characterized by "regulatory switching", where distinct lead variants or multiple molQTL types gained dominance at different stages. This is exemplified by *HSD17B12*, a key enzyme in fatty acid elongation^45,46^, which was controlled by two distinct, non-LD eQTL in early and peak lactation (PP.H4 = 0.03), suggesting a complete turnover of its regulatory drivers **(Fig. 6h)**.

To further dissect the mechanisms underlying this dynamic regulation, we next focused on post-transcriptional processes, particularly isoform usage ratio driven by alternative splicing, which can reshape protein outputs and ultimately influence phenotype, while largely uncoupled from gene-level expression changes^47,48^. We detected 549 stage-specific irQTL, of which 39 colocalized with stage-specific asQTL but only one colocalized with stage-specific sQTL, indicating that the application of lrRNA-seq substantially improved the resolution of regulatory effect mapping. A representative example is the peak-lactation signal for *AGPAT1* **(Fig. 6i).** At this locus, the variant rs211567677 was associated with the increased production efficiency of a specific isoform (XM_02498338.2) by modulating an alternative splicing event (**Fig. 6j**). Isoform usage gradually rise during lactation, peaking in late lactation (**Fig. 6k**). Given the role of *AGPAT1* in milk fat synthesis^49,50^, this case might illustrate how splicing-level variation dynamically fine-tunes switch major isoform to meet the high energetic demands of the mammary gland. Collectively, these results demonstrated that the bovine blood transcriptome undergoes systematic reprogramming of its regulatory architecture, reflecting a coordinated genetic response to the shifting immune, metabolic, and endocrine demands of lactation.

### Cellular composition and metabolic environments mediate lactation-specific regulatory effects

To determine whether cellular and metabolite shifts drive the stage-specific effects identified in the previous section, we systematically performed cell-type- and metabolite-interaction molQTL mapping across all 11 molecular phenotypes. Across seven major cell types, we identified 10,285 cell-type interaction molGenes (59.88% of all tested genes), while metabolite interaction analyses across 395 metabolites revealed 8,301 interaction molGenes (48.33%). Notably, lrRNA-seq data dramatically increased the resolution of interaction molQTL (iQTL) discovery. Compared with conventional eQTL analyses, isoQTL exhibited the highest sensitivity for detecting cell-type-dependent genetic effects (**Fig. 7a**). In contrast, metabolite interaction effects were most enriched at the splicing level, with sQTL showing the strongest signals (**Fig. 7a**). This layer-specific divergence suggests that cellular contexts primarily modulate allelic effects through isoform-expression regulation, whereas metabolic regulation is more directly related to functional splice variation might be due to their impacts on protein abundance or structure.

**Fig. 7.**
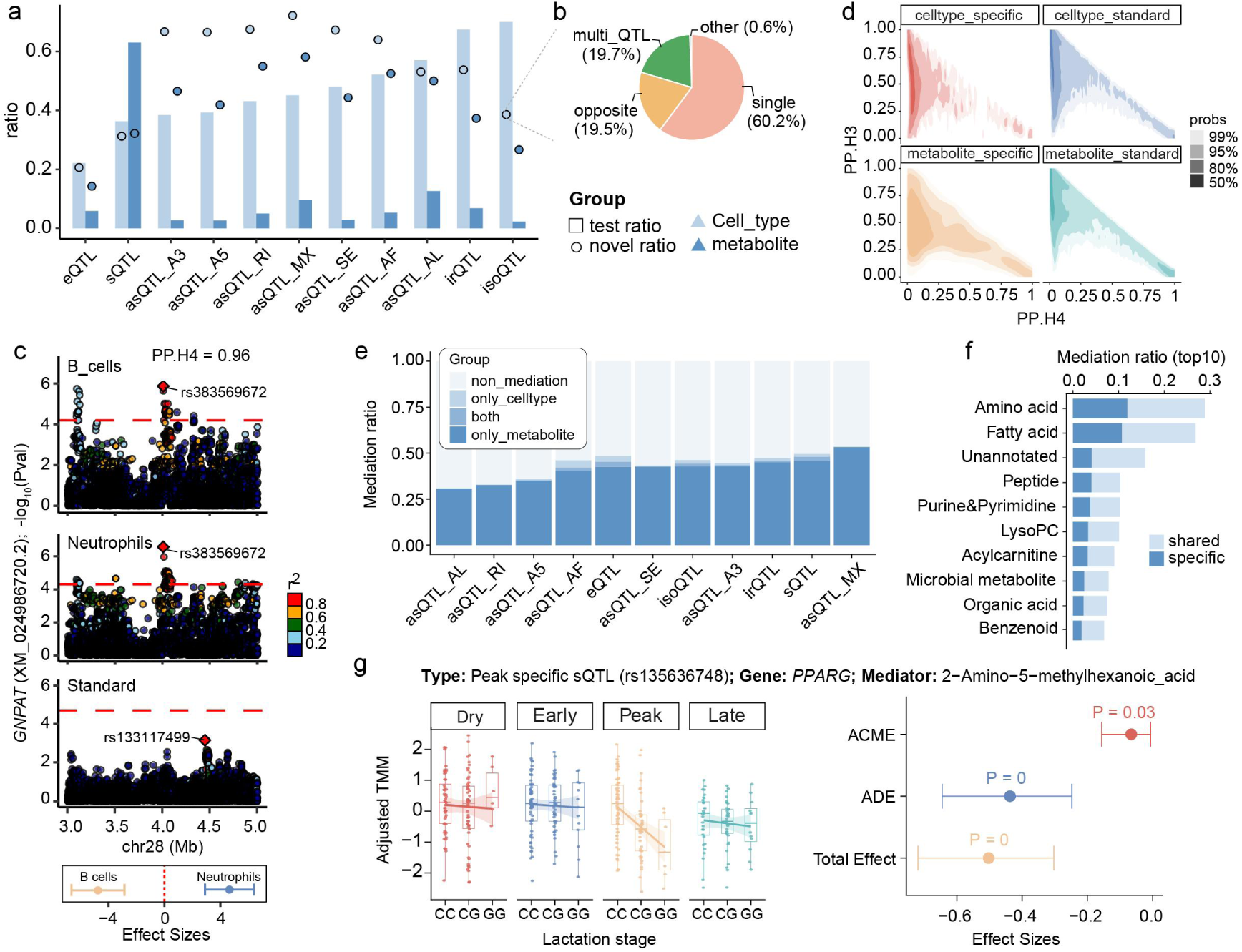
Mediation of lactation-stage-specific molQTL by cell type composition and metabolite abundance. **a,** Bar plot showed the detection efficiency of interaction molQTL mapping, and the scatter plot showed the proportion of novel molGenes in the interaction molGenes compared to the standard molGene. Light blue indicates cell type-interaction molGenes and dark blue indicates metabolite-interaction molGenes. **b,** Mechanistic categories underlying the identification of novel cell type-interaction isoGenes, including signals detected in only one cell type, signals detected in two cell types with opposite effect directions, signals shared across multiple cell types, and other categories. **c,** Representative example of a novel cell type-interaction isoGene showing significant interaction effects in two cell types with opposite directions, resulting in no detectable signal at the standard isoform level. **d,** Colocalization probabilities among interaction molQTL, lactation-stage-specific molQTL and standard molQTL. **e,** Proportion of lactation stage-specific molQTL mediated by cell type composition and metabolite abundance across different lactation stages. **f**, Proportion of lactation-stage-specific molQTL mediated by each metabolite class. molQTL mediated by metabolites from multiple classes were defined as shared. **g,** Representative example of a stage-specific molGene (*PPARG*) mediated by metabolite abundance.

Interaction molQTL mapping analyses uncovered a large proportion of cryptic regulatory effects that were completely masked in conventional bulk-level analyses. Relative to standard e/sQTL mapping based on srRNA-seq data, 20.7%-72.2% of cell-type interaction molGenes and 14.3%-58.1% of metabolite interaction molGenes were newly discovered (**Fig. 7a**). Notably, 38.6% of cell-type interaction isoGenes were novel. After excluding effects driven by rare cell populations, 19.5% of these isoforms displayed opposite allelic effects across different cell lineages, leading to complete signal cancellation in bulk-level (**Fig. 7b**). A representative example is the *GNPAT* isoform XM_024986720.2 showed strong colocalization between B-cell- and neutrophil-specific regulatory signals (PP.H4 = 0.96), but was undetectable in bulk analyses because the allelic effects acted in opposite directions across the two cell types (**Fig. 7c; Extended Data Fig. 6a**). These results demonstrate that a substantial fraction of the transcriptome is governed by antagonistic lineage-specific regulation that cannot be detected using conventional bulk-level approaches.

Despite abundant interaction signals, cell-type and metabolite interaction molPhenotypes showed limited explanatory power, accounting for only 39 and 45 stage-specific molPhenotypes, respectively, with merely 8 shared between the two categories **(Fig. 7d)**. This suggests a profound level of regulatory heterogeneity, where molecular effects often operate independently of cell-type and metabolite shifts (**Fig. 7c**). To further determine whether these contextual factors functionally transmit stage-specific regulatory effects, we performed mediation analysis using cellular composition and metabolite abundance as potential intermediates. Notably, metabolite abundance mediated 46.8% of stage-specific molecular phenotypes (**Fig. 7e**), far exceeding the contribution of cellular composition, which mediated only 4.3% of stage-specific effects **(Supplementary Table 9-10)**. Among metabolite-mediated signals, 41.4% were jointly mediated by multiple metabolite classes, with 96.2% classified as partial mediation and 3.9% as full mediation (**Extended Data Fig. 6b**). Amino acids and fatty acids were the dominant mediators, accounting for 28.9% and 26.9% respectively (**Fig. 7f**). Representative examples include peak-lactation-specific splicing of *PPARG*, a master regulator of adipogenesis and glucose homeostasis^51^, mediated by 2-Amino-5-methylhexanoic acid **(Fig. 7g)**, and dry-lactation-specific regulation of *EGR2* mediated by hexadecanedioic acid (FFA 16:0-DC) **(Extended Data Fig. 6c-e).** Collectively, these results demonstrate that while cell-type interactions provide high-resolution snapshots of lineage-specific regulation, metabolites act as the primary functional conduit through which genetic effects are transmitted to molecular phenotypes across the lactation cycle. This highlights the metabolic microenvironment as the dominant mediator of context-dependent gene regulation in dairy cattle.

### Regulatory architecture of GWAS loci and selection signatures in cattle

To dissect the regulatory basis of complex traits in cattle, we analyzed GWAS summary statistics from 39 metabolites and 105 complex traits, yielding 1,114 independent loci from 90 traits (**Supplementary Table 11**). Among all GWAS loci, asQTL showed the strongest enrichment (1.23-fold, s.e. = 0.017), followed by irQTL (1.21-fold, s.e. = 0.015), while eQTL and cell-type interaction eQTL showed lowest enrichment **(Fig. 8a, Extended Data Fig. 7a)**. Stage-specific molQTL also showed substantial enrichment **(Fig. 8b)**. Although metabolite iQTL (12.8%), eQTL (12.3%), sQTL(11.5%), and cell-type iQTL (11.0%) explained the largest proportion of *h²*, irQTL (2.07 × 10⁻⁵), asQTL (2.05 × 10⁻⁵), and isoQTL (1.49 × 10⁻⁵) exhibited the strongest per-variant effects on complex traits, indicating greater functional impact at individual loci despite lower overall contribution **(Fig. 8c; Supplementary Fig. 10)**.

**Fig. 8.**
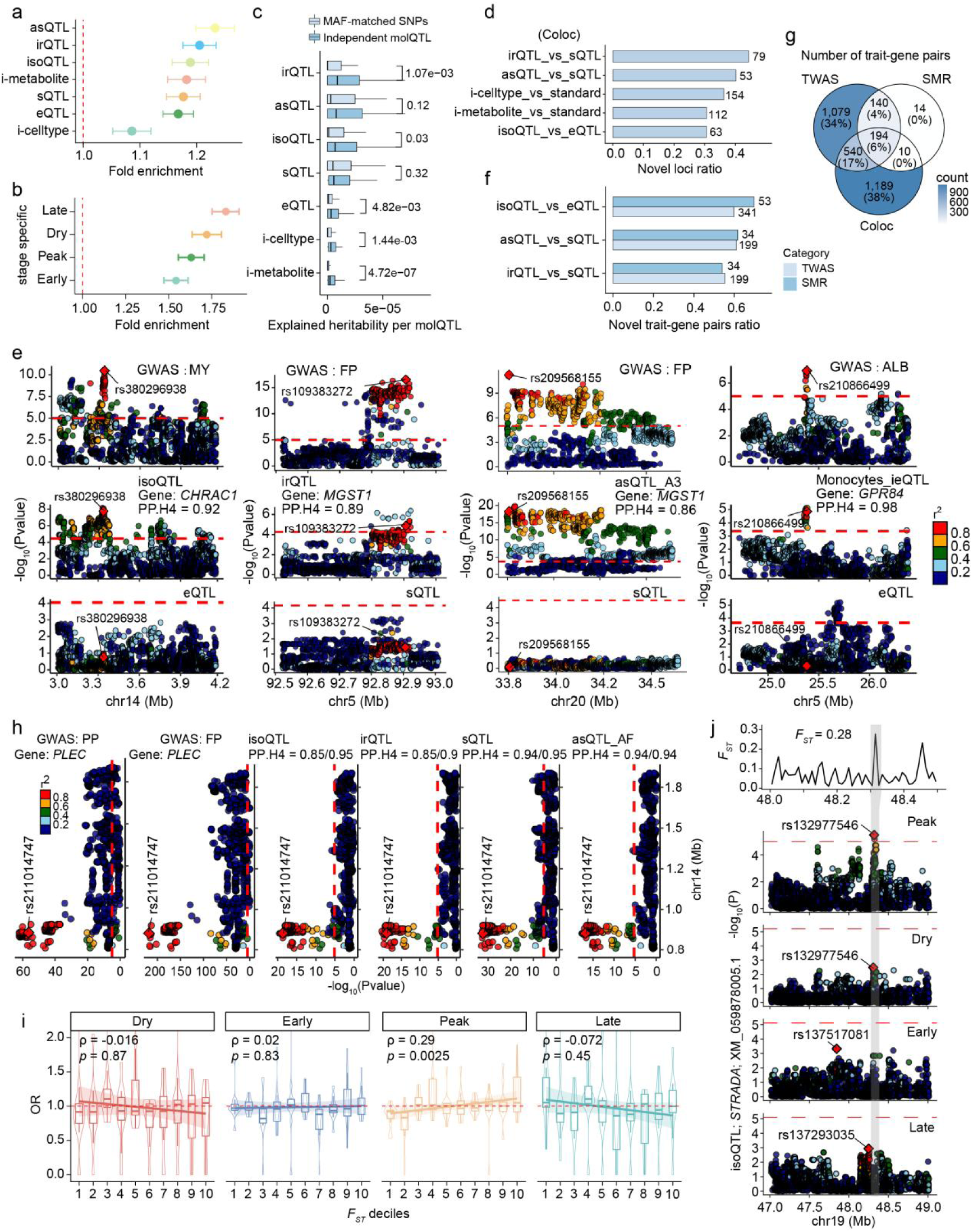
Interpreting GWAS loci of complex traits using molQTL. **a-b**, Enrichment (mean ± 95% confidence interval) of GWAS variants for 11 types of standard molQTL and interaction molQTL **(a)**, and lactation stage-specific molQTL **(b)**. **c**, Proportion of heritability explained for 90 cattle traits by independent molQTL versus MAF-matched random SNPs, correcting the number of molQTL. *P* values were calculated using two-sided paired Wilcoxon signed-rank tests. **d,** Number and proportion of GWAS-colocalized genes identified using transcript-level molecular phenotypes (isoQTL, irQTLA and asQTL) but not detected using gene-level molecular phenotypes (eQTL and sQTL). **e,** Examples of GWAS loci explained by only transcript-level and interaction molecular phenotype. **f,** Number and proportion of novel genes identified by SMR and TWAS using isoQTL relative to eQTL, and using asQTL or irQTL relative to sQTL. **g,** Venn diagram showing the overlap of trait-gene pairs identified by colocalization, TWAS and SMR. **h,** Example of *PLEC*, in which the same variant significant associated with protein percentage (PP) and fat percentage (FP) and explained by isoQTL, irQTL, sQTL, and asQTL_AF. **i**, Enrichment (odds ratio) of four classes of stage-specific molQTL across ten *F*_ST_ deciles. Linear regression lines indicate trends across selection intensity, and two-sided *P* values were obtained using Spearman correlation test. **j,** A representative peak-lactation-specific isoQTL under selection.

To associate molecular regulation with complex traits, we applied three complementary approaches including Bayesian colocalization (coloc)^52^, SMR^53^, and TWAS (S-PrediXcan)^54^. Overall, isoform-level molecular phenotypes together with interaction molQTL were colocalized with a large proportion of trait-associated variants missed by srRNA-seq analysis (eQTL and sQTL), increasing the proportion of explained GWAS loci from 35.2% to 60.3%, with the two layers contributing 5.7% and 19.5%, respectively, with 8.8% of the latter being long read length driven molecular phenotypes (**Extended Data Fig. 7b,c; Supplementary Table 12)**. More specifically, isoQTL colocalized with 206 GWAS loci, including 63 novel loci (5.7% of all GWAS loci) not captured by eQTL. Relative to sQTL, irQTL and asQTL colocalized with 79 (7.1%) and 53 (4.8%) novel GWAS loci, respectively **(Fig. 8d)**. Compared with steady molQTL, cell-type and metabolite iQTL colocalized with 154 (13.8%) and 112 (10.1%) novel GWAS loci, respectively. Representative examples included *CHRAC1* colocalizing with Milk Yield (MY) via an isoQTL (rs380296938), *MGST1* with Fat Percentage (FP) via an irQTL (rs109383272), *CARD6* with FP via an A3_asQTL (rs209568155), and *GPR84* with Albumin (ALB) via a monocyte ieQTL **(Fig. 8e)**. SMR and TWAS identified 358 and 1,953 significant trait-gene pairs, respectively, of which isoform-level analyses revealed 63 (54.3%) and 354 (46.9%) novel associations not detected at the gene level **(Fig. 8f; Extended Data Fig. 7d,e; Supplementary Table 13-14)**. Overall, three approaches converged on 194 high-confidence trait-gene pairs **(Fig. 8g)**. Representative example is the *PLEC* locus (rs211014747), associated with both PP and FP, which was simultaneously colocalized with isoQTL, irQTL, sQTL and AF_asQTL through the same lead variant **(Fig. 8h)** and identified by SMR and TWAS, yet exhibited opposite regulatory directions between splicing- and expression-based molQTL **(Supplementary Fig. 11)**, highlighting multilayer regulatory complexity. Similar examples include *FBXO24* for Body Volume (BV), *ADORA2A* for Dihydrouracil and Ureidopropionic acid, *MGST1* for FP, *DGAT1* for FP and PP, and *VPS28* for five production traits **(Extended Data Fig. 8)**.

To explore evolutionary forces shaping regulatory variation, we partitioned dairy-versus-beef cattle selection signals across *F*_ST_ bins and observed a strong positive association between selection intensity and peak-lactation-specific molQTL **(Fig. 8i)**, suggesting that domestication has preferentially targeted regulatory variants active during key physiological stages. For example, *STRADA* harbored a peak-lactation-specific isoQTL (rs132977546) within a highly selected region (*F_ST_* = 0.28) and encodes an important component of the mTOR signaling pathway, a central regulator of milk protein synthesis^40^ **(Fig. 8j)**. This supports a model in which milk production improvement is driven by coordinated remodeling of stage-specific regulatory networks.

Finally, to assess the contribution of molQTL to genomic prediction of 9 complex traits, including five milk producing traits, three body traits and one healthy trait, we incorporated molQTL into the GFBLUP genomic prediction framework in a Holstein population of 16,122. All molQTL explained 15.3%-87.0% of trait heritability **(Extended Data Fig. 9a)** and significantly improved prediction accuracy for milk yield (2.71%), fat percentage (9.14%), protein percentage (4.02%), and somatic cell score (3.77%), without compromising prediction unbiasedness **(Extended Data Fig. 9b,c; Supplementary Table 15)**. In contrast, MAF-matched random variants contributed minimal predictive power. These results support molQTL as a powerful functional layer for prioritizing molecular mechanisms of GWAS signals and enabling function-informed genomic selection.

### Cross-species conservation of regulatory variants between cattle and humans

To assess the cross-species conservation of bovine lead molQTL, we performed bidirectional liftOver to the human genome (hg38) with “-minMatch=0.5” and systematically evaluated their impacts on human genome function **(Fig. 9a)**. In total, 40.6% of molQTL (480,90 variants) were successfully mapped to orthologous human regions, with 30.3-52.9% conversion rates across molQTL types **(Fig. 9b)**. Using AlphaGenome^55^, orthologous bovine molQTL showed significantly higher predicted regulatory effects than MAF-matched random variants, indicating conserved regulatory function between humans and cattle **(Supplementary Table 16)**. Polytropic molQTL (QTL regulating at least two types of molQTL), asQTL, and sQTL displayed significantly higher predicted effect scores than eQTL, suggesting that splicing and multilayer regulatory variants are under stronger evolutionary constraint^56,57^ **(Fig. 9c)**.

**Fig. 9.**
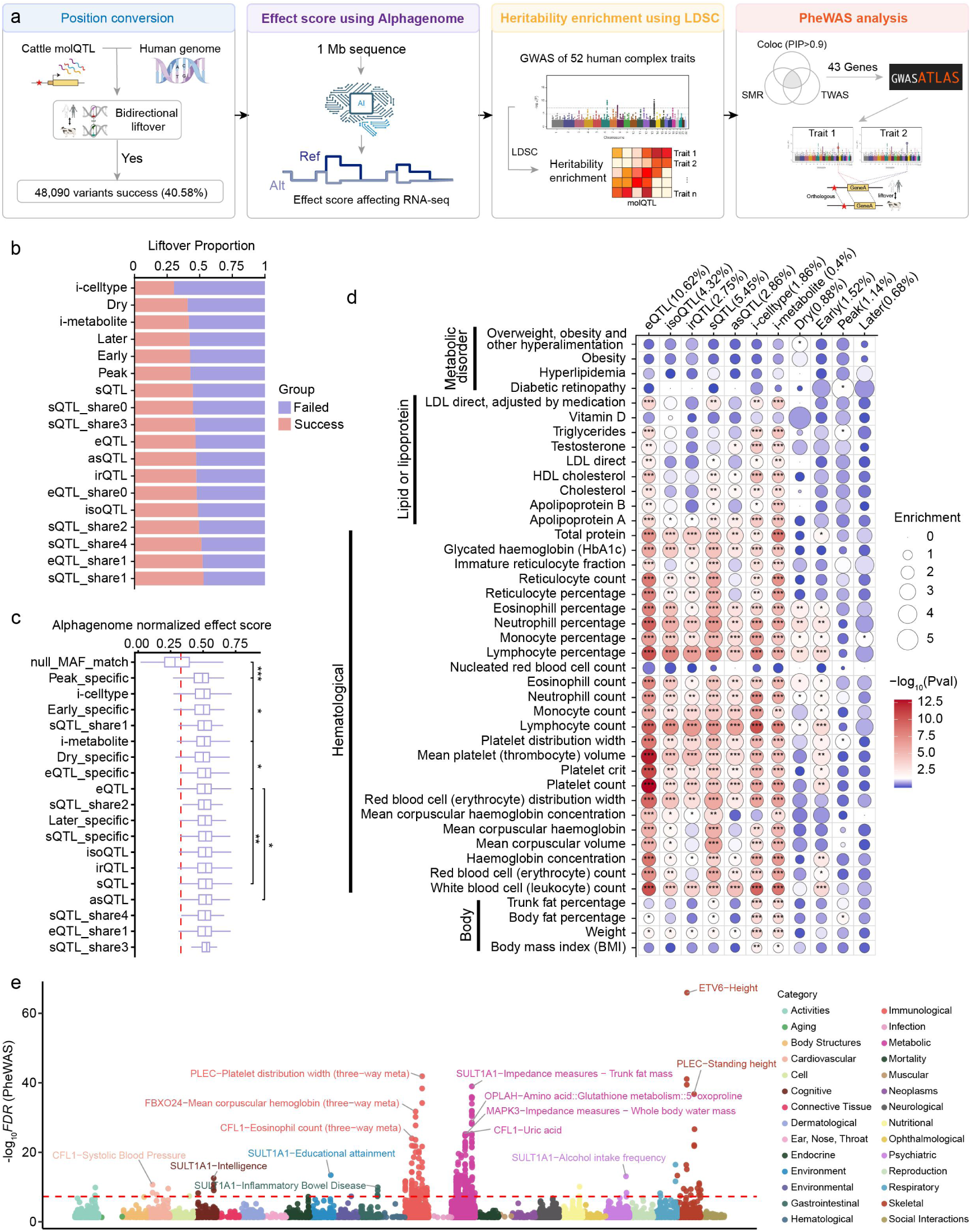
Evolutionary conservation of cattle molQTL in human complex traits. **a**, Overview of the analytical framework: (Ⅰ) reciprocal liftover of 11 types of standard lead molQTL, interaction molQTL, pleiotropic e/sQTL, lactation stage-specific molQTL, and MAF-matched random variants between the bovine and human genomes; (Ⅱ) AlphaGenome-based prediction of regulatory effects on human transcriptomes; (Ⅲ) partitioned heritability analysis via S-LDSC for 52 human complex traits; and (Ⅳ) PheWAS of cattle molQTL in human complex traits. **b**, Proportion of cattle molQTL successfully mapped to the human genome. **c**, AlphaGenome-predicted regulatory effects of liftover-mapped molQTL and MAF-matched random variants on human whole blood gene expression. Values are ordered from top to bottom by increasing mean effect. *P* values were calculated using two-sided paired Wilcoxon signed-rank tests. **d**, Partitioned heritability enrichment across 52 human complex traits. Numbers in parentheses indicate the proportion of SNPs covered by each molQTL. Dot size reflects the degree of enrichment, and color represents −log_10_(*P* value). **e**, Manhattan plot showing PheWAS results of cattle molQTL across human complex traits.

To further evaluate their contribution to human complex phenotypic variation, we applied stratified linkage disequilibrium score regression (S-LDSC) using orthologous bovine molQTL and GWAS summary statistics of 52 human complex traits **(Supplementary Table 17)**. Bovine molQTL showed significant heritability enrichment for hematological and metabolic traits, including apolipoprotein A, neutrophil percentage and lymphocyte count **(Fig. 9d)**. Notably, stage-specific regulatory variants showed distinct trait enrichments: dry- and early-specific molQTL were enriched for immune-related traits, whereas peak-specific molQTL were enriched for triglyceride, diabetic retinopathy, and body fat percentage traits, consistent with conserved physiological programs underlying lactation and metabolic remodeling. Furthermore, we performed a phenome-wide association study (PheWAS) across 4,756 human GWAS traits for 43 genes supported by colocalization, SMR, and TWAS evidence, using the criteria PP.H4 > 0.9, SMR FDR < 0.05, P_HEIDI > 0.05, and TWAS FDR < 0.05. In total, 15 genes showed significant associations with diverse human phenotypes (FDR < 5e-8; **Supplementary Table 18**), including *PLEC* with platelet distribution width, *SULT1A1* with trunk fat mass and inflammatory bowel disease (IBD), *DGAT1* with type 2 diabetes, *ETV6* with Height **(Fig. 9e)**. These findings highlight extensive evolutionary conservation of regulatory architectures and support the use of bovine functional genomics as a powerful comparative framework for understanding human complex traits.

## Discussion

Our study provides a multilayer framework for understanding how genetic variation propagates through transcriptomic regulation to shape complex phenotypes across dynamic physiological contexts in cattle. By integrating long- and short-read transcriptomics with genomics, metabolomics, and GWAS-scale phenotypic data throughout lactation, we systematically connected regulatory variation, transcript diversity, microenvironmental mediation, and organismal traits within a single population-scale resource. A central finding of this study is that a substantial fraction of regulatory variation operates beyond conventional gene-level expression^8,10,58^. Long-read transcriptomics enabled direct characterization of full-length isoforms and revealed widespread transcript-level regulatory effects that were undetectable in standard bulk analyses^18^. Many of these hidden signals arose from opposing, buffering, or dilution effects among isoforms, indicating that gene-level aggregation can obscure important components of functional genetic architecture. More broadly, the limited sharing of regulatory signals across molecular layers suggests that transcriptional regulation is not a simple linear hierarchy, but instead consists of partially independent modules governing expression abundance, isoform usage, and alternative splicing^7,8,10^. These observations support an emerging view that isoform-level regulation represents a major and previously underappreciated layer of complex trait biology.

Our results further demonstrated that genetic regulation is highly context dependent^59–61^. Regulatory effects varied markedly across lactation stages and were strongly shaped by cellular composition and metabolic environments. In many cases, regulatory variants exhibited opposite or compensatory effects across microenvironmental contexts, rendering their effects undetectable in conventional bulk analyses^59,61^. Interestingly, cellular composition preferentially influenced transcriptional regulation, whereas metabolites more strongly modulated functional splicing-related regulation, suggesting distinct mechanisms through which microenvironmental states reshape genetic effects. Together, these findings identify physiological context as a fundamental determinant of mammalian regulatory architecture and highlight the metabolic microenvironment as a major mediator of dynamic genetic regulation during lactation.

An important implication of this work is that integrating multiple regulatory layers substantially improves the interpretation of complex trait GWAS loci^62,63^. Transcript-level and context-dependent molQTL collectively resolved a large proportion of GWAS signals that remained unexplained by conventional eQTL-based approaches^63^. Moreover, many loci exhibited coordinated yet directionally discordant effects across multiple molecular phenotypes, emphasizing that regulatory variation frequently acts through interconnected but mechanistically distinct pathways. These findings underscore the importance of multi-omic integration for causal inference and suggest that a considerable fraction of the “missing regulation” underlying complex traits resides within isoform-level and context-specific regulatory mechanisms^63,64^.

Beyond mechanistic interpretation, our results also have important evolutionary and translational implications. Cross-species analyses revealed substantial conservation of regulatory architecture between cattle and humans, particularly for immune and metabolic traits, indicating that many context-dependent regulatory programs are conserved across mammals. These findings support the use of livestock functional genomics as a complementary framework for understanding human complex trait biology and disease mechanisms^6,10,65^. At the same time, incorporating functionally informed molQTL into genomic prediction models substantially improved predictive performance for economically important dairy traits, demonstrating the practical value of regulatory annotations for precision breeding.

Despite these advances, several limitations remain. Although our cohort represents one of the largest long-read multi-omics resources currently available in livestock, broader sampling across tissues, breeds, environments, and developmental stages will be necessary to fully capture regulatory diversity^20,66^. In addition, current long-read transcriptomic technologies still have limitations in sequencing depth and quantitative accuracy, particularly for low-abundance isoforms and complex splicing events^62,67,68^. Future single-cell long-read transcriptomic studies could resolve cell-type-specific isoform regulation that is masked in bulk tissues^69^, while direct lrRNA-seq may further enable joint profiling of full-length transcript structures and RNA modifications^70^. Furthermore, our analyses primarily focused on SNPs, whereas indels and structural variants, which may exert stronger effects on transcript structure and isoform regulation, remain largely unexplored. Therefore, future studies integrating larger genetically diverse cohorts from different breeds, multi-tissue profiling, longitudinal sampling, single-cell resolution, direct lrRNA-seq, and comprehensive structural variation analyses will further refine dynamic regulatory maps and deepen our understanding of mammalian gene regulation.

## Methods

### Ethics statement

The experimental procedures were approved by the Institutional Animal Care and Use Committee at the China Agricultural University (approval no. DK996).

### Sample and data collection

A total of 432 healthy primiparous Holstein dairy cows were randomly selected from a single commercial herd (Beijing Sunlon Livestock Development Co., Ltd., Beijing, China). Animals were stratified into four lactation stages, including dry (n = 112), early (n = 124), peak (n = 117), and late (n = 79) stages, with individuals within each stage have similar calving dates, body types, and feeding conditions **(Supplementary Table 1)**. Peripheral blood was collected from each cow by coccygeal venipuncture and used for downstream sequencing analyses.

### Whole-genome sequencing and variant calling

Genomic DNA was extracted from blood tissue using the DNeasy Blood & Tissue Kit (Qiagen, 69504) with RNase A treatment. DNA quality was evaluated using a NanoDrop spectrophotometer (Thermo Fisher Scientific) and verified by 1% agarose gel electrophoresis, and DNA concentration was quantified using a Qubit 2.0 fluorometer (Thermo Fisher Scientific). All samples were normalized to 40 ng/μL in 96-well plates. Sequencing libraries were constructed using the IGT® Enzyme Plus DNA Library Preparation Kit V3 (C11112, iGeneTech, Beijing, China) following the manufacturer’s standard protocol. Paired-end sequencing (2 × 150 bp) was performed on the DNBSEQ-T7 platform, achieving an average sequencing depth of 35.6×.

Raw reads were quality-filtered and adaptor-trimmed using fastp (v0.23.4)^71^. Clean reads were aligned to the ARS-UCD2.0 reference genome and generated individual GVCF using the “wgs” module in GTX.CAT (http://www.gtxlab.com/product/cat). Joint genotyping was performed by merging individual GVCF files with the “Joint” module of GTX.CAT. Variant calling was performed using GATK (v4.3.0), followed by variant selection with SelectVariants and quality filtering with VariantFiltration under the following criteria (QD < 2.0 || FS > 60.0 || MQ < 40.0 || MQRankSum < −12.5 || ReadPosRankSum < −8.0 || SOR > 3.0). After removing insertion variants, 12,937,265 high-quality variant were retained. For molQTL mapping, missing genotypes were imputed using Beagle (v5.4.0)^72^. SNPs were further filtered using PLINK (v1.9)^73^ by excluding variants with minor allele frequency (MAF) < 0.05, Hardy-Weinberg equilibrium P < 1 × 10^-6^, individual missing rate > 0.01, or genotype missing rate > 0.01. A total of 9,935,894 SNPs passed quality control and were used for downstream analyses.

### RNA extraction, library preparation, and sequencing

Total RNA was extracted from bovine blood using TRIzol (Thermo Fisher Scientific) and reverse-transcribed with the PrimeScript RT reagent Kit (Takara). RNA quality and concentration were assessed using NanoDrop and Agilent Bioanalyzer 2100. Strand-specific RNA-seq libraries were prepared and paired-end sequenced (150 bp) on the DNBSEQ-T7 platform. A total of 3.44 × 10^10^ raw reads were quality filtered and adapter trimmed using fastp (v0.23.4)^71^, resulting in a 3.43 × 10^10^ clean reads. These clean reads were aligned to the ARS-UCD2.0 reference genome (https://www.ncbi.nlm.nih.gov/datasets/genome/GCA_002263795.4/) using STAR (v2.7.9)^74^ with the parameters “--runMode alignReads --outSAMattributes NH HI AS nM XS --chimSegmentMin 10 --chimOutJunctionFormat 1 --alignSJoverhangMin 8 --twopassMode Basic --waspOutputMode SAMtag --outSAMtype BAM Unsorted --alignSJDBoverhangMin 1”. Aligned reads were sorted and deduplicated using samtools^75^.

Long-read cDNA libraries were prepared from 1 µg of total RNA using the cDNA-PCR Sequencing Kit (SQK-LSK110 + EXP-PCB096) provided by Oxford Nanopore Technologies (ONT). Full-length cDNA was generated using template-switching reverse transcription, followed by PCR amplification (14 cycles) with LongAmp Tag (Ipswich, MA, USA) and adaptor ligation using T4 DNA ligase. Libraries were purified with Agencourt XP beads (Beckman, Indianapolis, IN, USA) and sequenced on FLO-MIN109 flow cells on the PromethION platform. Raw electrical signal data were basecalled using dorado (https://github.com/nanoporetech/dorado/). Full-length reads were identified using Pychopper with primer configuration “+:SSP, −VNP | −:VNP, −SSP”, where VNP and SSP corresponded to ACTTGCCTGTCGCTCTATCTTC and TTTCTGTTGGTGCTGATATTGCT, respectively. Reads shorter than 100 bp or with a mean quality score < 7 were removed using chopper (v0.8.0)^76^ with parameters “-q 7 −l 100”, yielding 4.68×10^9^ post-QC reads and 4.25×10^12^ bases. Clean reads were aligned to the ARS-UCD2.0 reference genome using minimap2 (v2.28)^77^ with parameters “-ax splice -ub -k14”, followed by sorting and deduplication with samtools^75^. Sequencing quality was assessed using NanoPlot (v1.43.0)^76^.

### Non-targeted Metabolomic sequencing

Serum samples were analyzed by ultrahigh-performance liquid chromatography coupled with high-resolution mass spectrometry (UPLC-HRMS) for untargeted metabolomics profiling. Serum proteins were precipitated using ice-cold methanol:acetonitrile (1:3, vol/vol) followed by centrifugation (15,000 × g, 2 min, 4°C). The resulting supernatant was lyophilized (−50°C, 10 Pa) and reconstituted in 25% methanol (vol/vol) for analysis. A pooled quality control (QC) sample was generated by combining 10 μL of each extract and injected every ten samples to monitor the stability and reproducibility of the UPLC–HRMS system. Polar metabolites were annotated by comparison with in-house chemical standards (Sigma-Aldrich, St. Louis, MO), a local SMOL spectral library (Beijing Bioyun Huakang Genomics Co. Ltd, Beijing, China), HMDB v5.0 (Human Metabolome Database, https://hmdb.ca) and the mzCloud spectral database (Thermo Scientific, https://www.mzcloud.org/). Raw data files were converted to mzXML format and processed using XCMS, CAMERA and metaX in R (v4.2.0). Molecular features were defined by retention time (RT) and mass-to-charge ratio (m/z), and peak intensities were extracted to generate a three-dimensional data matrix comprising RT–m/z features, sample identities and ion intensities. Metabolite identification and quantification were performed by Beijing BioMiao Biological Technology (Beijing, China).

### Hybrid transcriptome assembly and quantification

To balance sensitivity and specificity in transcript discovery, we used a hybrid assembly strategy that integrates short-read and long-read RNA-seq and requires concordance across multiple assembly algorithms. In brief, short- and long-read transcriptome data were jointly integrated to perform hybrid transcriptome assembly for each sample using FLAIR (v2.0.0)^26^, StringTie2 (v2.2.2)^25^, and RNA-Bloom, separately (v2.0.1)^27^. For each assembly approach, isoforms were merged across samples using StringTie2 (v2.2.2)^25^, and isoforms supported by at least two of the three assembly approaches were retained using gffcompare (v0.12.6)^78^. These retained isoforms were subsequently merged to produce a unified transcriptome annotation using StringTie2. Transcript abundance was then quantified using Salmon (v1.10.3)^79^ based on this annotation. Assembled transcripts were further curated, refined, and classified using SQANTI3 (v5.2) with default parameters^28^. Isoforms novelty was assessed by comparison with the ARS-UCD2.0 reference annotation, resulting in the identification of 13,177 novel isoforms. These novel isoforms were merged with the ARS-UCD2.0 reference annotation to generate a blood-enhanced transcriptome annotation comprising 94,712 transcripts. Open reading frames (ORFs) of novel isoforms were predicted using TransDecoder (v5.7.1; https://github.com/TransDecoder/TransDecoder). Predicted peptide sequences were aligned to the Swiss-Prot database using DIAMOND (v2.1.9)^80^, and protein domains were annotated using hmmsearch against the Pfam database. The protein-coding potential of novel isoforms was further assessed using CPC2^30^, CNCI^31^, and CPAT (v3.0.5)^32^. Isoforms predicted to lack coding potential by at least two methods were classified as non-coding transcripts, whereas genes harboring at least one protein-coding isoform were defined as protein-coding genes. DIAMOND was also used to infer the potential biological processes associated with novel isoforms based on COG database (https://ftp.ncbi.nih.gov/pub/COG/KOG/).

### Quantification of molecular phenotypes

Clean RNA-seq reads were aligned to the ARS-UCD2.0 reference genome with the blood-enhanced transcriptome atlas. Isoform expression was quantified using the quant module of Salmon (v1.10.3)^79^, and gene-level counts were summarized using tximport (v1.3.9)^81^, with expression normalized to transcripts per million (TPM). Genes and isoforms with reads raw count < 6 or TPM < 0.1 in more than 20% of samples were filtered, resulting in 17,176 expressed genes and 40,224 expressed transcripts. Remaining read counts were normalized using trimmed mean of M values (TMM), followed by the inverse normal transformation. Isoform ratio was calculated as the TPM of each isoform divided by the sum of TPM values of all isoforms of the same gene, yielding 33,451 isoform ratio phenotypes, which were further processed by quantile normalization followed by inverse normal transformation.

Alternative splicing was quantified using Leafcutter (v0.2.7)^82^ on WASP-filtered alignments. Splice junction reads were extracted using the bam2junc.sh script, and intron clusters were defined using leafcutter_cluster.py (--min_clu_reads 30 --min_clu_ratio 0.001 --max_intron_len 500000), and mapped to genes using the map_clusters_to_genes.R. Introns were filtered according to established criteria^6,83^, excluding those with > 50% missing or zero counts, fewer than max(10, 0.1n) unique values (n = sample size), or low complexity (∑_i_(|z_i_| < 0.25) ≥ n − 3 and ∑_i_(|z_i_| > 6) ≤ 3, where z_i_ is the z score of the *i*th cluster read fraction across individuals). Remaining counts were normalized using prepare_phenotype_table.py, yielding 86,780 alternative splicing phenotypes.

Alternative splicing events were identified using SUPPA (v2.3)^84^. All possible splicing events were generated from the blood-enhanced transcriptome atlas using the generateEvents function, and quantified at the local splicing and isoform levels using psiPerEvent and psiPerIsoform, respectively. Initially identified 7,645 skipped exon (SE), 6,680 alternative first exon (AF), 5,414 alternative 3′ splice site (A3), 4,854 alternative 5′ splice site (A5), 2,091 retained intron (RI), 1,455 alternative last exon (AL), and 888 mutually exclusive exon (MX) events. For each category, events with valid PSI values in at least 80% of individuals were retained, resulting in 7,481 SE, 6,435 AF, 5,247 A3, 4,701 A5, 2,061 RI, 1,414 AL, and 855 MX events, which were subsequently quantile-normalized and inverse normal transformation for downstream analysis. For all molecular phenotypes, missing values were imputed using a k-nearest neighbors (KNN) approach, and molecular phenotypes with zero variance across individuals were removed prior to molQTL mapping.

### Metabolome data analysis

Metabolite data were preprocessed and normalized were performed using MetaboAnalystR^85^. Metabolites with missing values in all samples or low variability (based on interquartile range and mean abundance) were removed. The filtered data were normalized using total sum scaling, log transformation, and autoscaling. Seven outlier individuals were identified based on the OPLS-DA results excluded from downstream analyses.

A genomic relationship matrix (GRM) was constructed using GCTA (v1.93)^86^ with the --grm option, and genomic principal components (PCs) were estimated using --pca. Narrow-sense heritability of each metabolite was estimated using restricted maximum likelihood (REML), with lactation stage fitted as a fixed effect and the top three genomic PCs included as covariates. Genome-wide association analyses were performed using a mixed linear model in GCTA (--mlma)^87^. Genome-wide significance threshold was set at P < 9.37×10^-9^, based on Bonferroni correction for 410,688 independent variants and 13 effective independent metabolites. Independent genetic variants were identified using PLINK (--indep-pairwise 50 5 0.2). The effective number of independent metabolites was estimated by eigenvalue decomposition of the metabolite covariance matrix, accounting for correlations among metabolites, using the formula Neff = (∑λ)² / ∑(λ²). Independent association signals were further identified using conditional and joint analysis with GCTA-COJO^88^ (--cojo-p 9.37 × 10⁻⁹ --cojo-wind 1000 --cojo-collinear 0.9 --cojo-slc).

To identify marker metabolites used to distinguish dry and lactating, normalized metabolomics data were analyzed using a repeated machine learning framework implemented in the caret (v7.0.1; https://github.com/topepo/caret/) package. Samples were classified into lactating vs dry. Seven supervised classifiers (glmnet, LogitBoost, partial least squares, regularized logistic regression, Random Forest, linear SVM, XGBoost) were trained using 10-fold cross-validation. For XGBoost, these parameters were fixed (nrounds = 100, max_depth = 3, eta = 0.1, colsample_bytree = 0.7, subsample = 0.7). In each of 10 repetitions, 25 samples per group were randomly selected as an independent test set, while the remaining samples were split into training and validation sets at a 9:1 ratio using a fixed random seed. Model performance was evaluated on the test set using ROC curves and AUC values computed with pROC (v1.19.0.1)^89^, and the optimal model was selected based on the highest mean AUC across repetitions. The selected model was then retrained on the full dataset and interpreted using model-agnostic KernelSHAP. SHAP values were computed as class probabilities for the target group, and metabolite importance was quantified by mean absolute SHAP values (global importance) and signed SHAP values within the target group (directional contribution). SHAP-based feature importance, distribution (beeswarm), and individual prediction explanations were visualized using shapviz (v0.10.3)^42^.

### Differential expression, co-expression network, and time-series analyses

Genes with TPM > 0.1 in at least 20% of samples were retained, and the corresponding raw count matrix was used for downstream analyses. Hidden batch effects were inferred and removed using SVA (v3.5.8)^90^ package, with lactation stage specified as the variable of interest (model: ∼ lactation stages; null model: ∼ 1), resulting in 36 surrogate variables and corrected expression matrices for downstream analyses. Differential expression analysis was performed using DESeq2 (v1.50.2)^91^ with surrogate variables included as covariates (design formula: ∼ SV1 + SV2 + lactation stages) to identify differentially expressed genes (DEGs) across lactation stages. Differential abundence metabolites (DAMs) were identified from preprocessed metabolite abundances using limma (v3.66.0)^92^, with P values adjusted by the Benjamini-Hochberg (BH) procedure. Genes or metabolites with false discovery rate (FDR) < 0.05 and an absolute log_2_ fold change (|log_2_FC|) > 0.58 were considered significantly differentially.

Gene co-expression networks were constructed separately for each lactation stages using normalized TMM data with WGCNA (v1.71)^93^. Pearson correlation all between all gene pairs were calculated to generate a correlation matrix. An appropriate soft-thresholding power (β) was determined using the scale-free topology criterion. Adjacency matrices were then used to construct gene networks and identify modules via a one-step dynamic tree cutting algorithm. Module eigengenes (first principal component of module expression) were tested for association with stages, and modules with *P* < 0.05 were considered candidate co-expression modules. Hub genes were defined as those with gene significance (GS) > 0.2 and module membership (MM) > 0.8.

For time-series analysis across lactation stages, gene expression values were transformed as log_2_(CPM + 1), and metabolite abundances were log_2_-transformed after normalization. Values were averaged within each lactation stage before clustering. The optimal number of clusters was determined using the elbow method. Temporal expression and abundance patterns were identified using fuzzy c-means clustering implemented in Mfuzz (v2.70.0)^94^, and cluster-specific trajectories were visualized using ClusterGVis (v0.99.5, https://github.com/junjunlab/ClusterGVis).

### Cell type deconvolution of bulk blood transcriptomes

Based on a previously constructed multi-tissue single-cell transcriptomic atlas of cattle^41^, 46,078 blood cells were classified into 13 distinct cell types. Cell type-specific expression signatures were generated with the buildSignatureMatrixMAST function in DWLS (v1.82.0)^95^. Bulk blood gene expression profiles (TPM) were subsequently deconvolved using solveDampenedWLS function to estimate the relative proportions of each cell type for each individual. Cell types with significant differences in predicted proportions across lactation stages and an average proportion > 2% were retained, resulting in seven cell types for downstream analyses.

### cis-molQTL mapping

Standard cis-molQTL mapping was performed using OMIGA (v1.1.3)^96^ with a mixed linear model accounting for the additive polygenic background effect. Cis-windows were defined as ±1 Mb around the TSS of the each gene. Briefly, nominal associations for all variant-molecular phenotype pairs were computed using the “--mode cis --qtl-map-model a+A” option. PCs from molecular phenotypes and genotypes were automatically inferred, with the optimal number determined by a convergence criterion (--dprop-pc-covar 0.001). Highly collinear covariates with an absolute correlation coefficient greater than 0.95 were removed (--rm-collinear-covar 0.95), and lactation stage was included as a fixed-effect covariate using the --covariates option. For cis-isoQTL, cis-irQTL, cis-sQTL, and cis-asQTL analyses, molecular phenotypes were grouped by gene using the --pheno-group option, and gene-level multiple testing correction was performed using Clipper(--multiple-testing clipper). Variants with nominal *P* values (pval_g1) below the corresponding significance threshold (pval_g1_threshold) were defined as molQTL, and genes with adjusted q values (qval_g1) < 0.05 harboring at least one molQTL were designated as molGenes. *Cis-h*^2^ of molecular phenotypes was estimated using OMIGA (--mode her_est --h2_model Ac), and independent cis-molQTL signals were identified using stepwise conditional mapping (--mode cis_independent). For isoQTL and irQTL, genes containing only one isoform are excluded.

Interaction molQTL (iQTL) analyses were conducted to evaluate the influence of cellular composition and metabolic abundance on genetic regulation. Interaction terms were incorporated using the “--interaction” option with estimated cellular proportions or metabolite abundances as interaction variables. Cis-interaction molQTL mapping was performed using the “--mode cis_interaction” option. For cell-type interaction molQTL mapping, the mixed linear model was retained. Variants with pval_g2 below the corresponding significance threshold were defined as cell-type iQTL, and genes with qval_g2 < 0.05 harboring at least one iQTL were designated as molGenes. For metabolite interaction QTL, a general linear model (--qtl-map-model a) was applied due to reduced detection power, and multiple testing correction was performed using the aggregated Cauchy association test (ACAT; --multiple-testing acat). Variants with pval_g2_acat < 0.05 were defined as metabolite iQTL, and genes with qval_g2 < 0.05 were considered molGenes.

### Fine-mapping of molQTL

Putative causal variants for molGenes were fine-mapped using the “Sum of Single Effects” (SuSiE) model (v.1.0)^97^ in TensorQTL (v1.0.9)^98^. Summary statistics for the cis-region of each Phenotype were used as input, while the corresponding genotype and linkage disequilibrium (LD) matrix were generated using PLINK (v1.9)^73^. Variants with posterior inclusion probabilitie (PIP) summing up to 95% or higher were identified as credible sets (CS).

### Functional annotation and enrichment analysis of molQTL

Fine-mapped variants for molGenes were annotated and tested for enrichment using OMIGA (--mode enrich --enrich-annot). Variants in credible sets were annotated with functional genomic features and 10 chromatin states, and enrichment was assessed relative to a background set of MAF-matched random variants to control for MAF-dependent biases. Functional consequences of variants were annotated using SnpEff (v5.4)^99^. Chromatin state annotations were obtained from a cattle epigenome atlas^35^. Chromatin state abbreviations: TssA, Active TSS; TssFlnk, Flanking TSS; TssWk, Weak (Flanking) TSS; EnhA, Active Enhancer; EnhWk, Weak Enhancer; EnhPos, Poised Enhancer; BivTss, Bivalent TSS; BivEnh, Bivalent Enhancer; ReprPC, Repressed Polycomb; Quies, Quiescent/low.

### Structural impact of molQTL variants on proteins

Fine-mapped variants were classified according to protein sequence effects, predicted structural effects and posterior inclusion probability (PIP). Four variant sets were defined: set1, MAF-matched background variants with low confidence (PIP < 0.5); set2, high-confidence non-protein-sequence-altering variants (PIP > 0.8); set3, high-confidence protein-altering variants (PIP > 0.8); and set 4, high-confidence protein-altering variants (PIP > 0.8) with predicted structural effects. Protein-altering fine-mapped variants located within coding sequences were further evaluated for their effects on protein sequence and structure. Variant alleles were normalized against the reference genome using bcftools (v1.23)^100^ “norm”, and sample-specific sequences were generated using “bcftools consensus”. Coding sequences and translated protein sequences were extracted using gffread (v0.12.8)^101^. For variants altering amino acid sequences, reference- and alternative-allele protein sequences were generated and their structures were predicted using AlphaFold3^102^. Structural differences were quantified using global root-mean-square deviation (RMSD), maximum predicted aligned error (PAE) change, and Cα atom distance change. Variants were considered to affect protein structure if they met any of the following criteria: global RMSD > 2 Å, maximum PAE change > 5, or Cα distance change > 1 Å.

### Shared and specific molQTL across lactation stages

Standard cis-molQTL mapping was first performed for 11 molecular phenotypes at each lactation stage, followed by a cross-lactation stage meta-analysis using MashR (v0.2.79)^103^. For each molecular phenotype, z-scores (effect size divided by standard error) derived from OMIGA (v1.1.3)^96^ were extracted for the lead cis-molQTL and used as input for MashR. MashR estimated posterior effect sizes and statistical significance, quantified as local false sign rates (LFSR). A molQTL was defined stage-specific if it had LFSR < 0.05 in only one stage. Genetic similarity of gene expression regulation across stages, was assessed by Spearman correlation of posterior effect sizes estimates for variants with LFSR < 0.05 in at least one group. A molGene was defined as stage-specific if it harbored at least one stage-specific molQTL.

### Mediation analysis

To assess whether lactation stage-specific molQTL regulatory effects are mediated by cellular composition or metabolic abundances, mediation analyses were performed using the mediation (v4.5.0)^104^ package. Cellular composition and metabolite abundances were treated as potential mediators, and stage-specific molecular phenotypes were used as outcomes. For each variant–phenotype–mediator triplet, We fitted one mediator model and one outcome model:

Mediator model:

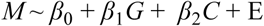

Outcome model:

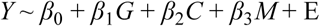

where *Y* represents molecular phenotype, *M* denotes the mediator, *G* is the genotype, *C* is the covariates used for QTL mapping, *β*_1_ is the genotype effect, *β*_2_denotes the covariate effect, *β*_3_is the mediator effect, and E is residuals. The significance of the Average Causal Mediation Effects (ACME), Average Direct Effects (ADE), Total Effects and Proportion of Mediated Effects was evaluated at a 95% confidence using nonparametric bootstrap (N = 999 Monte Carlo draws). Mediated effects were considered significant if ACME *P* < 0.05.

### GWAS summary statistics of complex traits

To investigate the regulatory mechanisms underlying complex traits in cattle, identified molQTL were integrated with GWAS summary statistics for 90 complex traits in Holstein cattle, including 23 production traits, 18 health traits, 39 metabolite traits, four body conformation traits, five reproductive traits, and one longevity trait. For all traits, a genomic relationship matrix was constructed using GCTA (v1.93) with the --grm option, and genomic principal components were estimated using --pca. Narrow-sense heritability was estimated by REML, with lactation stage fitted as a fixed effect and the top three genomic principal components included as covariates. GWAS was then performed using a mixed linear model in GCTA (--mlma)^87^. Before downstream integration, GWAS and molQTL summary statistics were harmonized to the same reference genome, SNP identifiers, effect alleles and genomic coordinates. Ambiguous or unmatched variants were removed, and allelic effects were flipped when necessary to ensure consistent effect directions across datasets.

### Enrichment of molQTL and trait-associated variants

To assess enrichment of molQTL among trait-associated variants, two complementary approaches were used. First, QTLEnrich (v2)^5^ was applied to quantify enrichment between significant molQTL and GWAS loci across 90 bovine complex traits. Second, variants were partitioned into molQTL, MAF-matched SNPs, and remaining SNPs. Category-specific GRMs were constructed, and REML-based variance component analysis in GCTA was used to estimate the proportion of phenotypic variance explained by each SNP category.

### Colocalization analysis

Trait-associated independent SNPs were identified from GWAS summary statistics using GCTA-COJO with the options --cojo-p 1×10^-6^, --cojo-wind 1000, --cojo-collinear 0.9 and --cojo-slc, using ∼16,000 Holstein cattle as the LD reference panel. For metabolite traits, the COJO significance threshold was set to 9.37×10^-9^. For each independent lead SNP, the corresponding GWAS locus was defined as the ±1 Mb region around the lead variant.

To identify shared genetic variants between GWAS and cis-molQTL, colocalization analysis were performed for each molGene and overlapping GWAS loci using the coloc.abf function in the coloc (v.5.2.3)^105^ package. This Bayesian framework evaluates five hypotheses: (1) no association with either GWAS loci and molQTL (H0); (2) association only with GWAS loci (H1); (3) association only with molQTL (H2); (4) association with both GWAS loci and molQTL but two independent signals (H3); and (5) association with both GWAS loci and molQTL driven by a shared causal variant (H4). Pairwise LD between lead GWAS loci and molQTL was calculated using PLINK. To ensure robust colocalization inference, GWAS locus-molPhenotype pairs with fewer than 500 shared SNPs were excluded from downstream analysis.

### TWAS analysis

Transcriptome-wide association study (TWAS) were performed using S-PrediXcan implemented in MetaXcan (v.0.8.0)^54^. For each of the 11 molecular phenotypes, gene expression prediction models were trained using nested cross-validated elastic net regression with cis-SNPs within ±1 Mb of each gene. The predictive models with cross-validated correlation ρ > 0.1 and prediction P < 0.05 were retained. These models were then applied to GWAS summary statistics using S-PrediXcan to infer gene-trait associations at the molecular phenotype levels. Significant associations were defined as those with FDR< 0.05.

### SMR analysis

SMR analysis was performed to assess the association between genetically regulated molecular phenotypes and bovine complex traits. Cis-molQTL summary statistics were used as exposures, and GWAS summary statistics of 90 traits were used as outcomes. For each molPhenotype, the top cis-variant was used as the instrumental variant, and LD was estimated from the WGS cohort. The HEIDI test was applied to distinguish pleiotropic associations from linkage. Significant SMR associations (FDR < 0.05) without HEIDI heterogeneity (*P* > 0.05) were retained as candidate trait-associated regulatory molecular phenotypes.

### Enrichment of molQTL in selection signature

Selective sweeps were inferred by contrasting 651 dairy (Holstein) and 552 beef (326 Simmental, 64 Angus, 60 Charolais, 58 Limousin and 44 Hereford) cattle using genome-wide SNPs. Genomic windows were ranked by *F*st and stratified into ten deciles representing increasing levels of population differentiation. Enrichment of molQTL within each Fst decile was evaluated using odds ratios (OR), defined as

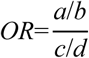

Where *a* is the numbers of molQTL in a given *F*st decile, *b* is the total number of molQTL, *c* is the number of SNPs in that decile, *d* is the number of all SNPs. Allele frequencies were calculated using the option of “--freq” in PLINK.

### molQTL-driven GFBLUP

Genomic prediction was evaluated in a reference population of 16,122 Holstein cattle, including 15,636 individuals for milk yield (MY), protein yield (PY), fat yield (FY), protein percentage (PP), fat percentage (FP) and somatic cell score (SCS) trait, and 11,709 individuals for conformation (CONF), mammary system score (MS) and foot & leg (FL). Three models were compared using hiblup (v1.6.0)^106^: standard GBLUP, molQTL-informed GFBLUP, and GFBLUP based on randomly selected MAF-matched variants, allowing assessment of the predictive contribution of molQTL. All models were evaluated using 10-fold cross-validation procedure with five repetitions.

GBLUP model:

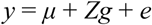

GFBLUP model:

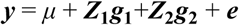

where ***y*** is the vector of trait deregressed proof (DRP), *μ* is the overall mean, ***g*** is the vector of direct genomic value (DGV), ***e*** is the vector of random errors, and ***Z*** is an incidence matrix allocating records to ***g***. The random effects follow a normal distribution 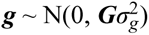 and 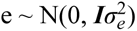, where **G** is the genomic matrix, and 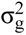 and 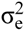 are the additive genetic variance and the residual variance, respectively. In the GFBLUP model, *g*_1_ and *g*_2_ represent the DGV captured by functional variants (e.g., molQTL and random selected SNPs) and the remaining variants, respectively, with their corresponding GRM constructed separately.

Prediction accuracy was measured by the correlation between the DGV and DRP, and can be calculated via 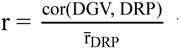 where r̅**_DRP_** is the average DRP accuracy. Prediction unbiasedness was evaluated as the regression coefficient of DRP on DGV, and can be calculated via 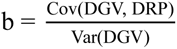 with b=1 indicating unbiased prediction.

### Cross-species comparison between cattle molGenes and human complex traits

To assess the evolutionary conservation of cattle molGenes in human complex traits, three complementary analysis were perfomed. First, lead molQTL of each molecular phenotype were mapped from the bovine genome to the human reference genome (hg38) using reciprocal liftover^107^ (http://hgdownload.cse.ucsc.edu/hubs/GCF/002/263/795/GCF_002263795.3/liftOver/GCF_002263795.3ToHg38.over.chain.gz; http://hgdownload.cse.ucsc.edu/goldenPath/hg38/liftOver/hg38ToGCF_002263795.3.over.chain.gz). For each mapped variant, reference and alternative alleles were obtained from dbSNP v153 (http://fileserve.mrcieu.ac.uk/dbsnp/dbsnp.v153.hg38.vcf.gz), and regulatory effects in human Whole_Blood were predicted using AlphaGenome (v0.6.1)^55^. A 1-Mb window (±500 kb) centered on each variant was scored using the model.score_variant function, restricted to RNA-SEQ tracks from whole blood. Quantile-normalized absolute effect sizes were averaged to derive a final functional impact score for each molQTL.

Second, GWAS summary statistics for 52 complex traits from the UK Biobank were used to tested whether human orthologs of molQTL were enriched for trait heritability using LD score regression (LDSC, v1.0.1)^108^. Liftover-mapped molQTL were treated as a functional annotation, and partitioned heritability was estimated as the proportion of SNP-based heritability explained by molQTL relative to their proportion of total SNPs.

Final, the relevance of cattle molGenes to human complex traits by phenome-wide association analysis (PheWAS) using the GWASATLAS database (https://atlas.ctglab.nl/PheWAS), which includes 4,756 GWAS summary statistics covering 4,285 human traits. There are a total of 43 molGenes supported by colocalization, SMR, and TWAS evidence, using the criteria PP.H4 > 0.9, SMR FDR < 0.05, P_HEIDI > 0.05, and TWAS FDR < 0.05. Subsequently, PhWAS analysis was performed on their human orthologs.

### Functional Enrichment Analysis

Gene Ontology (GO) and KEGG pathway enrichment analyses were performed using clusterProfiler (v4.0)^109^. GO terms and KEGG pathways were tested using the org.Bt.eg.db and bta databases with the erichGO and enrichKEGG functions, respectively, with a threshold of *P* < 0.05, visualized using ggplot2 (v3.5.2) and aPEAR (v1.0.0)^110^.

### Data availability

The raw WGS, long- and short-read RNA-seq, and metabolome data newly generated in this study can be retrieved from the Sequence Read Archive under accession numbers PRJCA062649 at Nation Genomics Data Center (https://ngdc.cncb.ac.cn). The ARS-UCD2.0 cattle reference genome is available at NCBI (https://www.ncbi.nlm.nih.gov/datasets/genome/GCF_002263795.3/). All GWAS and molQTL summary statistics results can be found in Supplementary Tables and were available to download at https://cattleblr.farmgtex.org/download.

### Code availability

All the computational scripts and codes for WGS, long- and short-read RNA-seq, and metabolome analyses, as well as the respective quality control, molecular phenotype normalization, genotype imputation, hybrid transcriptome assembly, molQTL mapping, functional enrichment, colocalization, TWAS and selection sweep are available at the GitHub website (https://github.com/bioramen-Blip/Long-read-Blood-cattleGTEx-Pipeline).

## Acknowledgements

This work was financially supported by the National Key R&D Program of China (2021YFF1000700); High Innovation Plan (202504841083); STI 2030-Major Projects (2023ZD04069); the Program for Changjiang Scholar and Innovation Research Team in University (IRT_15R62); the 2115 Talent Development Program of China Agricultural University. Lingzhao Fang was supported by Agriculture and Food Research Initiative Competitive grants nos. 2022-67015-36215 from the USDA National Institute of Food and Agriculture, seed-funding from CellFood Hub (AUFF), Danmarks Frie Forskningsfond nos. 5254-00069B. Jianguo Li was supported by Top Talent Project of Hebei Province (6012018). We acknowledge the support of the High-Performance Computing Platform of China Agricultural University (Beijing) and the Xihe High-Performance Computing Platform of the National Research Facility for Phenotypic and Genotypic Analysis of Model Animals (Beijing).

## Author information

These authors jointly supervised this work: Dongxiao Sun, Lingzhao Fang.

## Authors and Affiliations

**State Key Laboratory of Animal Biotech Breeding, National Engineering Laboratory for Animal Breeding, Key Laboratory of Animal Genetics, Breeding and Reproduction of Ministry of Agriculture and Rural Affairs, College of Animal Science and Technology, China Agricultural University, Beijing 100193, China.**

Weijie Zheng, Qi Zhang, Jinfeng He, Zijiao Guo, Yanan Liu, Aixia Du, Xiaoning Zhu, Bo Han, Yu Zhang, Dongxiao Sun.

**Center for Quantitative Genetics and Genomics, Aarhus University, Aarhus 8000, Denmark.**

Mian Gong, Houcheng Li, Bingjin Lin, Bingxing An, Di Zhu, Xinfeng Liu, Lingzhao Fang.

**Beijing Dairy Cattle Center, Beijing 100192, China.**

Lin Liu, Zhu Ma.

**Institute of Animal Science and Veterinary Medicine, Shandong Academy of Agricultural Sciences, Jinan, 250000, China.**

Jianbin Li.

**Key Laboratory of Genetic Evolution & Animal Models, Kunming Institute of Zoology, Chinese Academy of Sciences, Kunming, Yunnan 650201, China**

Jingsheng Lu, Xuemei Lu.

**Department of Animal Science and Technology, Hebei Agricultural University, Baoding, 071001, China**

Jianguo Li.

## Author contributions

All authors made substantial contributions to the conception or design of the study; the acquisition, analysis or interpretation of data; or drafting or revising the paper. D. Sun and L. Fang conceived and designed the project. D. Sun, L. Liu, J. Li, Z. Ma, and J. Li provided samples and data. W. Zheng performed all bioinformatic analyses. Q. Zhang provided assistance with transcript assembly. L. Liu, J. Li, Z. Ma, J. He, Z. Guo, Y. Liu, A. Du and B. Han helped collect samples and complex trait data of dairy cows, as well as GWAS analysis. X. Zhu, M. Gong, H. Li, Y. Zhang, B. Lin, B. An, D. Zhu, and X. Liu provided assistance with molQTL mapping analysis. W. Zheng, J. Lu, and X. Lu jointly completed the website construction. D. Sun contributed to the data and computational resources. L. Fang and D. Sun contributed to the critical interpretation of analytical results before and during manuscript preparation. W. Zheng drafted the manuscript. L. Fang and D. Sun revised the manuscript. All authors read, edited and approved the final manuscript.

## Competing interests

The authors declare no competing interests.

## Extended Data Figures

**Extended Data Fig. 1.**
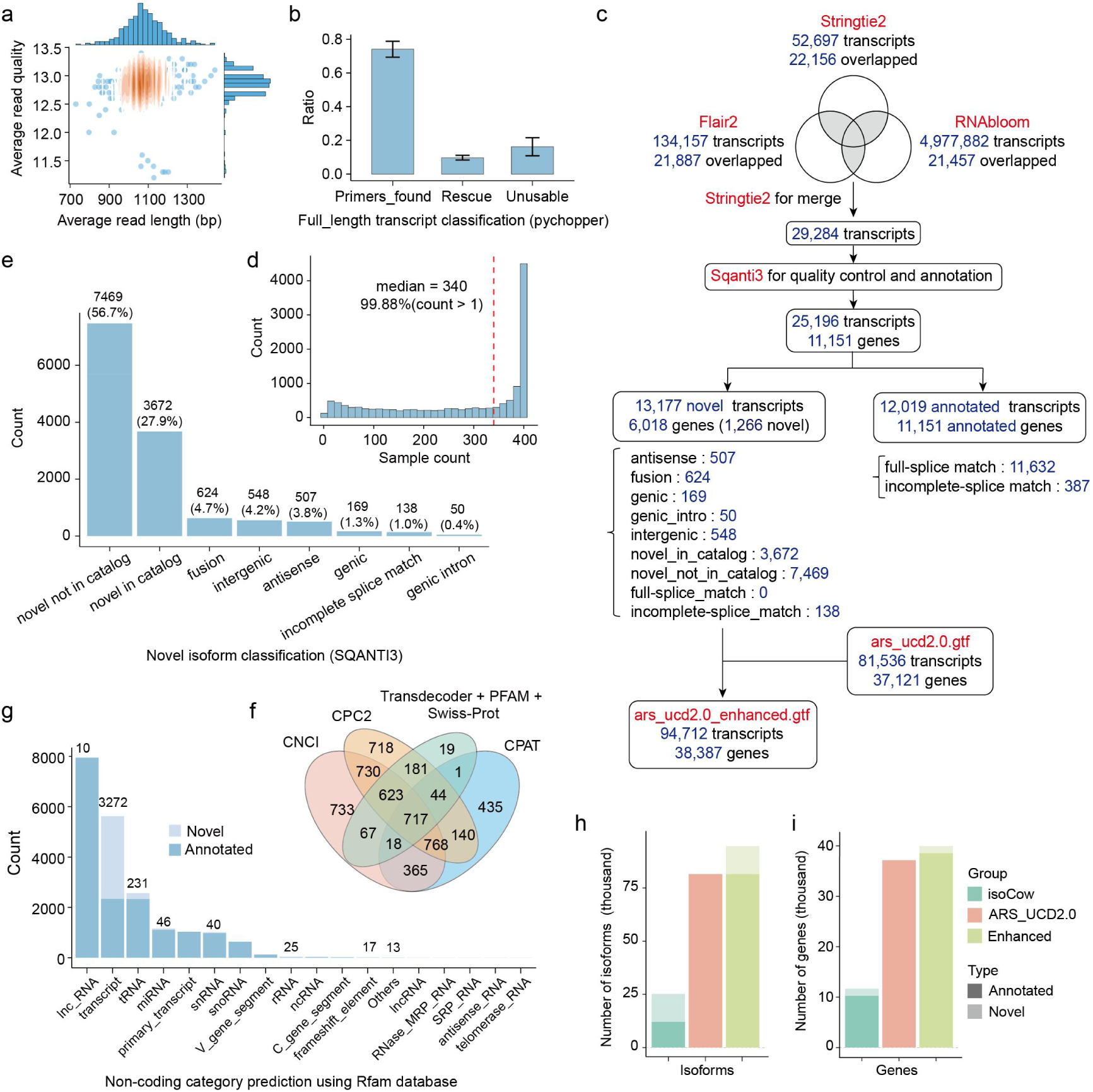
Construction and classification profiling of an enhanced transcriptome annotation. **a,** Joint distribution of average read length and average read quality across all RNA-seq libraries. **b,** Proportion of full-length reads identified by Pychopper, rescued full-length reads and non-full-length reads from long-read transcriptome sequencing. **c,** Workflow for transcriptome assembly and annotation. **d,** Distribution of the number of samples in which novel isoforms was detected by long-read RNA-seq. The red dashed line indicates the median sample count per isoform (median = 340), and 99.88% of isoforms were detected in more than one sample. **e,** Classification of novel isoforms by SQANTI3. **f**, Venn diagram showing the overlap of protein-coding potential predictions from CPC2, CNCI, CPAT and TransDecoder combined with Pfam and Swiss-Prot searches, used to define non-coding isoforms. **g**, Distribution of non-coding categories of isoforms. **h-i,** Comparison of isoform **(h)** and gene **(i)** count among isoCow, ARS-UCD2.0 and the enhanced atlas, showing annotated and novel isoform proportions.

**Extended Data Fig. 2.**
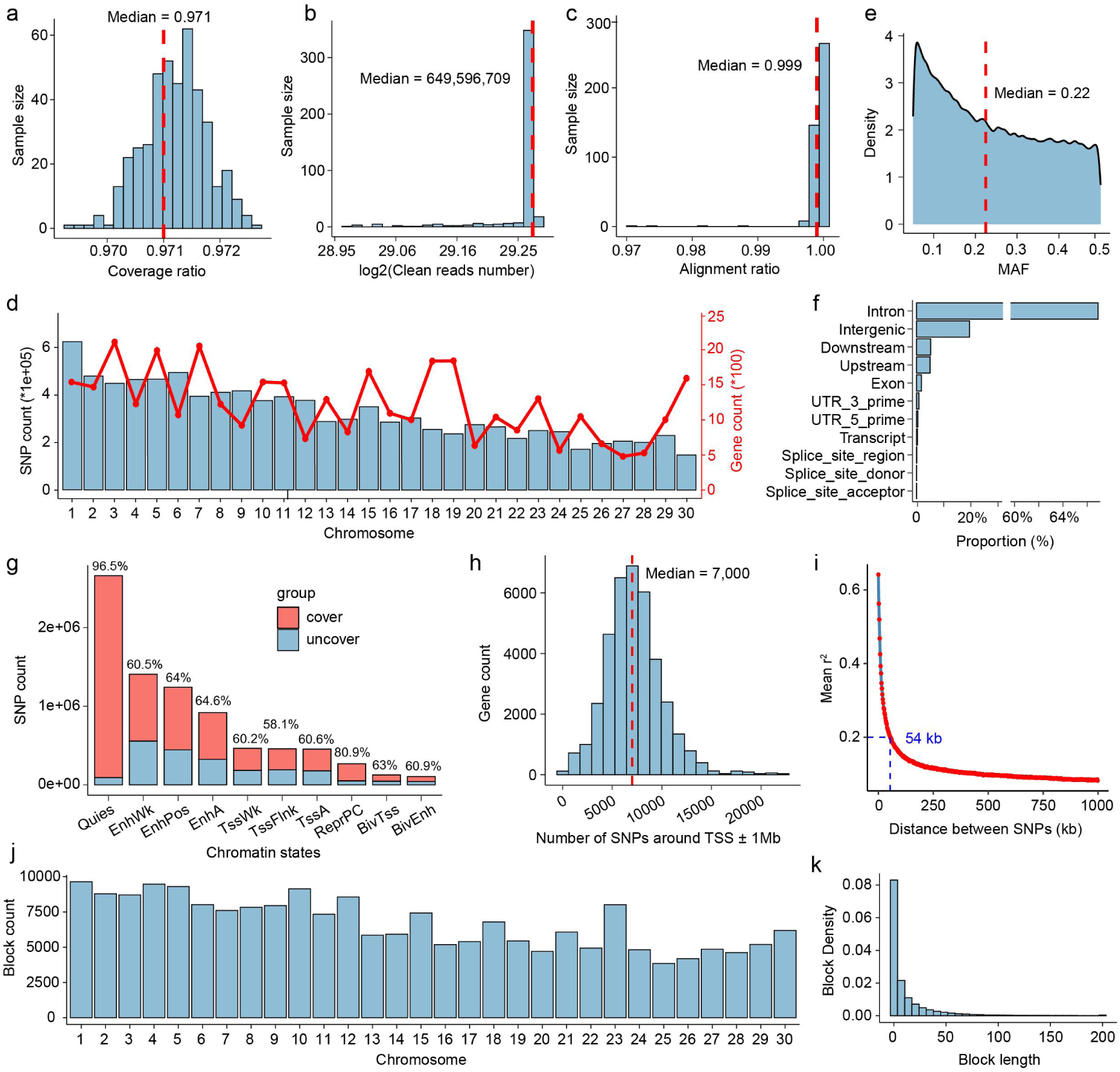
Data summary of WGS. **a,** Distribution of genome coverage ratio across all samples, with a median of 0.971. **b**, Distribution of clean sequencing depth shown as log2-transformed number of clean reads (median = 649,596,709). **c**, Alignment ratio of clean reads to the reference genome, showing a median of 0.999. **d**, Chromosome-wise distribution of SNPs (grey bars) and genes (red line). **e**, Minor allele frequency (MAF) distribution of all SNPs, with a median of 0.22. **f**, Functional annotation of SNPs, showing their genomic locations relative to gene features, including intronic, intergenic, exonic and regulatory regions. **g,** Distribution of genomic regions with or without SNP coverage across ten chromatin states. Red indicates regions covered by at least one SNP, whereas blue indicates regions without SNP coverage. **h,** Distribution of the number of SNPs located within ±1 Mb of transcription start sites (TSSs), with a median of 7,000 SNPs per gene. **i,** Linkage disequilibrium (LD) decay across the cattle genome. Pairwise LD decreased to *r²* < 0.2 at an inter-SNP distance of ∼54 kb. **j,** Number of LD blocks identified on each chromosome. **k,** Distribution of LD block lengths, with a median length of 1.8 kb.

**Extended Data Fig. 3.**
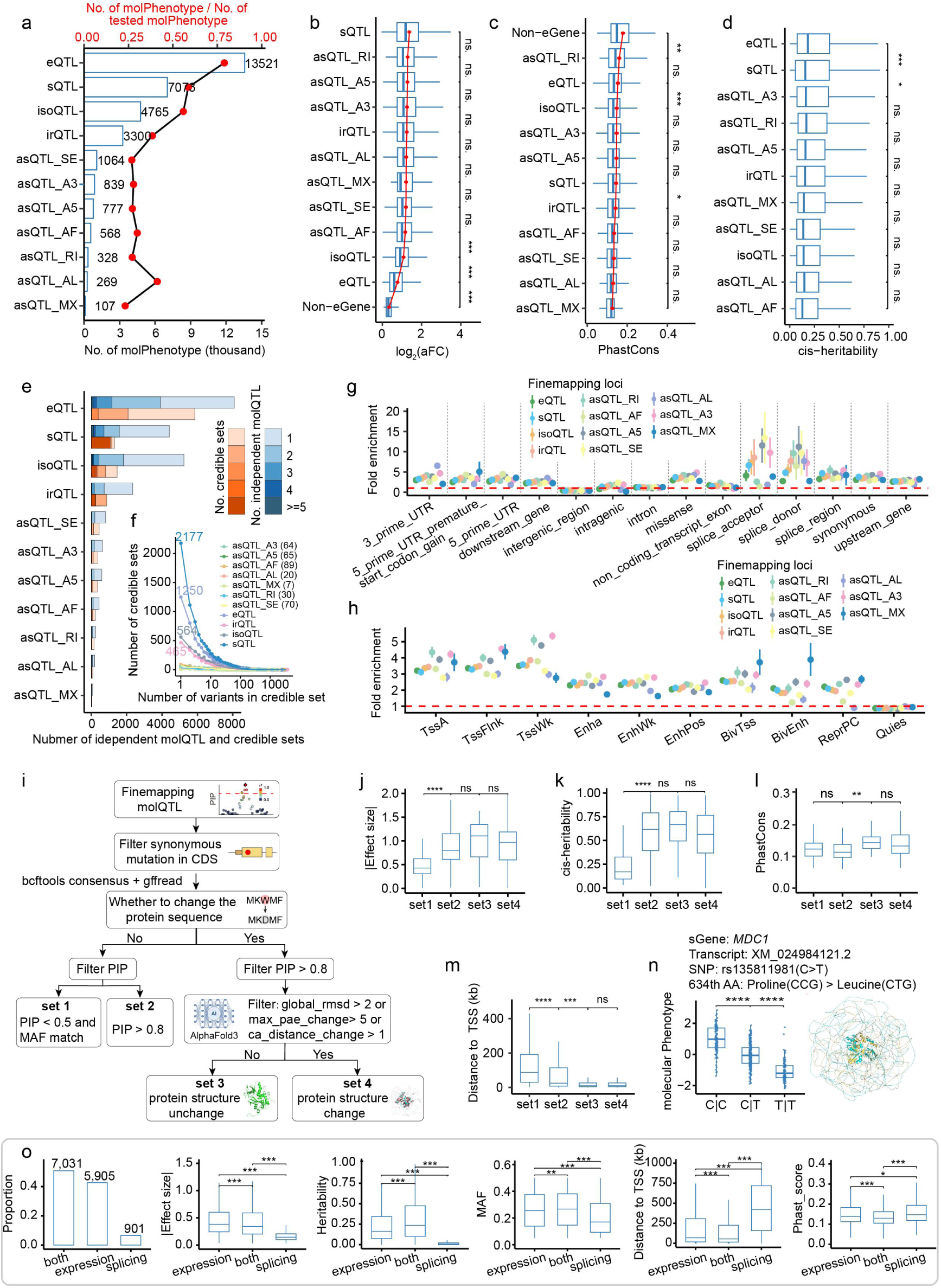
Genetic and functional analysis of molQTL across regulatory layers. **a,** Numbers of molGenes detected for each of 11 molecular phenotypes and corresponding detection efficiencies. **b-d**, Box plots comparing the allelic Fold Change (aFC) **(b)**, PhastCons score **(c)**, and *cis-h2* **(d)** across molQTL. *P* values were calculated using two-sided paired Wilcoxon signed-rank tests. *P* values were calculated using two-sided paired Wilcoxon signed-rank tests. NS: non-significant; *: *P* < 0.05; **: *P* < 0.01; ***: *P* < 0.001. **e,** Number of independent molQTLs and fine-mapped credible sets across the 11 molecular phenotypes. **f,** Distribution of credible-set sizes across molecular phenotypes. Values indicate the number of credible sets containing a single candidate causal variant. **g-h,** Enrichment of fine-mapped molQTL variants across 14 sequence ontology categories **(g)** and 10 chromatin states **(h)**. Points indicate mean enrichment estimates and error bars indicate 95% confidence intervals. **i,** Fine-mapping and structural-impact workflow, stratifying variants by PIP and protein-structure changes. All fine-mapped variants were classified into four categories: set 1, non-protein-altering variants with low PIP (< 0.5) matched for MAF; set 2, non-protein-altering variants with high PIP (> 0.8); set 3, protein-altering variants with high PIP predicted not to affect protein structure; and set 4, protein-altering variants with high PIP predicted to alter protein structure. **j-m**, Box plots comparing the |β| **(j)**, *cis-h*^2^ **(k)**, PhastCons scores **(l)**, and distance to TSS **(m)**. *P* values were calculated using two-sided paired Wilcoxon signed-rank tests. NS: non-significant; *: *P* < 0.05; **: *P* < 0.01; ***: *P* < 0.001. **n,** Example of a protein-altering sQTL. Box plot shows genotype-dependent expression differences. Right panel shows superimposed three-dimensional protein structures before and after the amino-acid substitution. **o,** Proportion, |β|, *cis-h*^2^, MAF, distance to the TSS, and PhastCons conservation score of molGenes regulated by expression-based molQTL, splicing-based molQTL or both. *P* values were calculated using two-sided paired Wilcoxon signed-rank tests. NS: non-significant; *: *P* < 0.05; **: *P* < 0.01; ***: *P* < 0.001.

**Extended Data Fig. 4.**
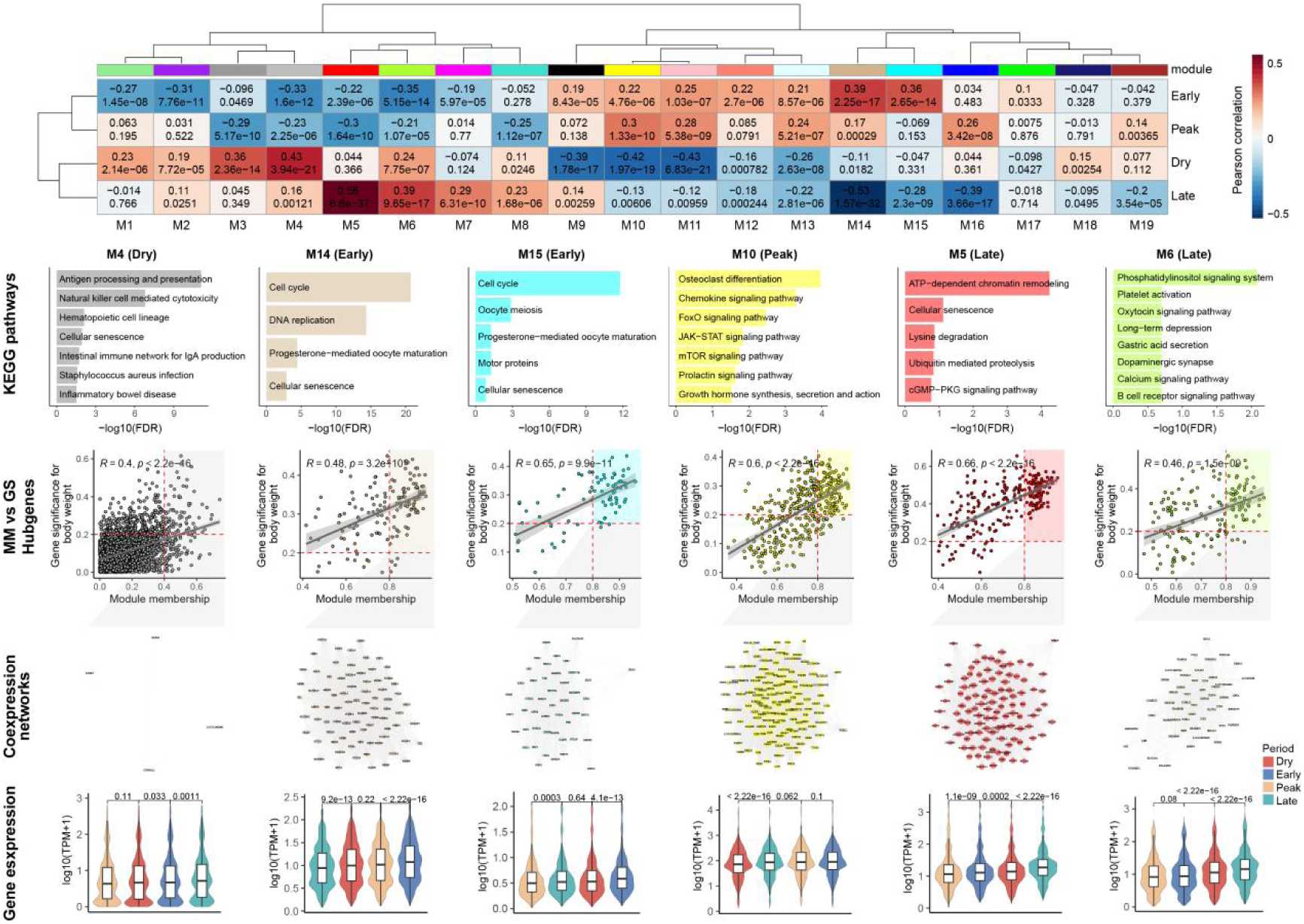
Gene co-expression network analysis across lactation stages. From top to bottom, heatmap plot shows 19 gene co-expression modules significantly associated with the four lactation stages; Bar plot shows highlights KEGG pathway enrichment of module genes significantly associated with each stage, including M4 (dry), M14 and M15 (early), M10 (peak), M5 and M6 (late); Scatter plot displays the distribution of gene significance (GS) versus module membership (MM) for each module, with linear trends fitted using geom_smooth from the ggplot2 package. Two-sided Pearson correlation tests were used to assess significance. Hub genes were defined as GS > 0.2 and MM > 0.8 (MM > 0.4 for M4), and co-expression networks of module genes were visualized using igraph package; Box plot shows stage-specific expression of hub genes across lactation stages. *P* values were calculated using two-sided paired Wilcoxon signed-rank tests.

**Extended Data Fig. 5.**
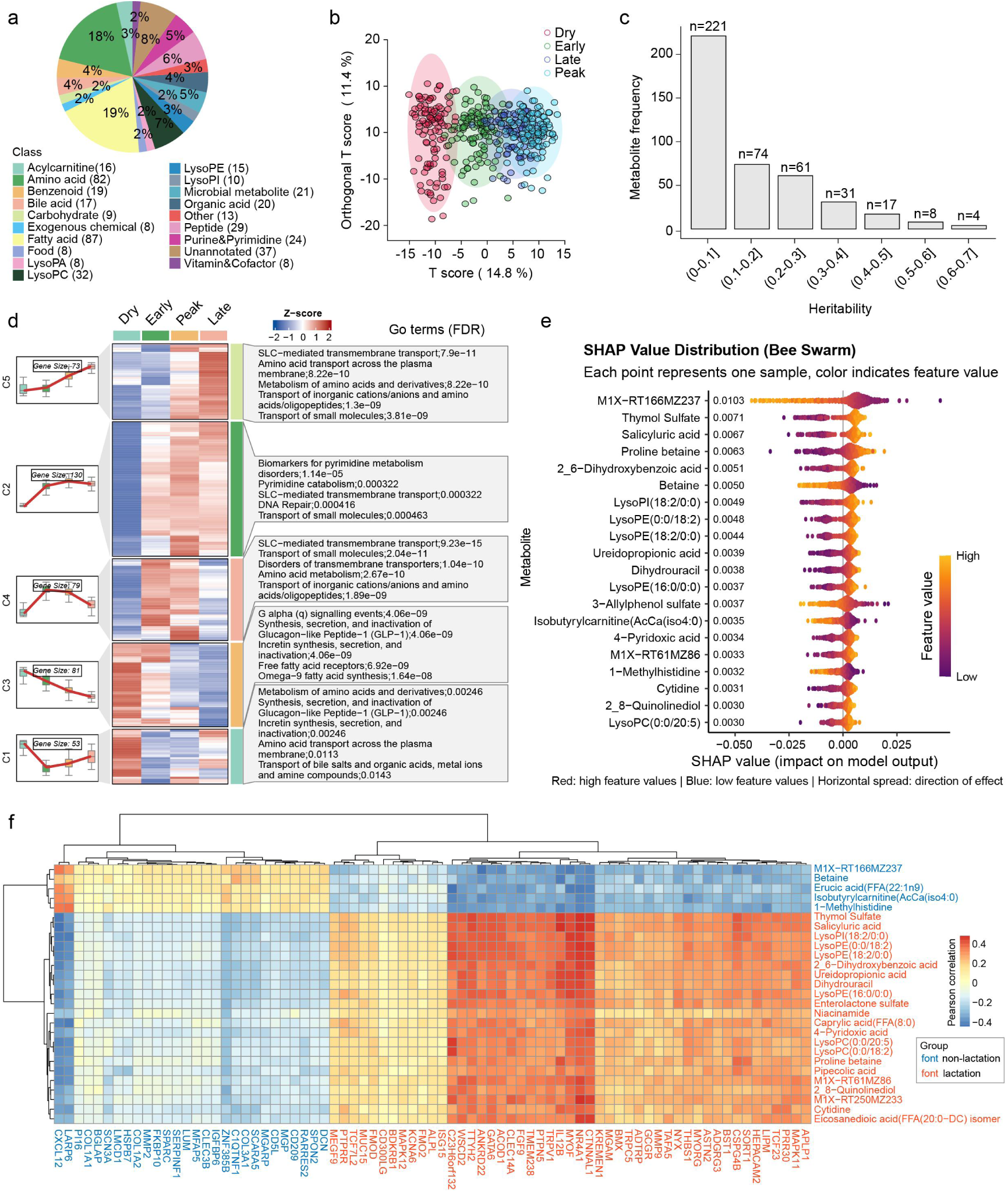
Metabolic reprogramming during lactation. **a**, Pie chart showing the distribution of 19 metabolite classes, with fatty acids and amino acids comprising the largest proportions, 87 metabolites (19%) and 82 metabolites (18%), respectively. **b**, OPLS-DA analysis based on metabolite abundance reveals clear clustering of individuals at each of the four lactation stages. **c**, Bar plot illustrating the number of metabolites grouped by heritability, with a step size of 0.1. **d**, Time-series analysis identified five distinct expression trends. Heatmap plot shows the abundance of different modules across the four lactation stages, with gene-enriched GO terms and their corresponding FDR values on the right. **e**, SHAP values showing the contribution of individual metabolites to model predictions; the top 20 most informative metabolites are highlighted. **f**, Heatmap showing correlations between differentially expressed genes and differentially abundant metabolites in dry versus lactation comparisons. Blue labels indicate genes and metabolites upregulated in the dry stage, and red labels indicate those upregulated during lactation.

**Extended Data Fig. 6.**
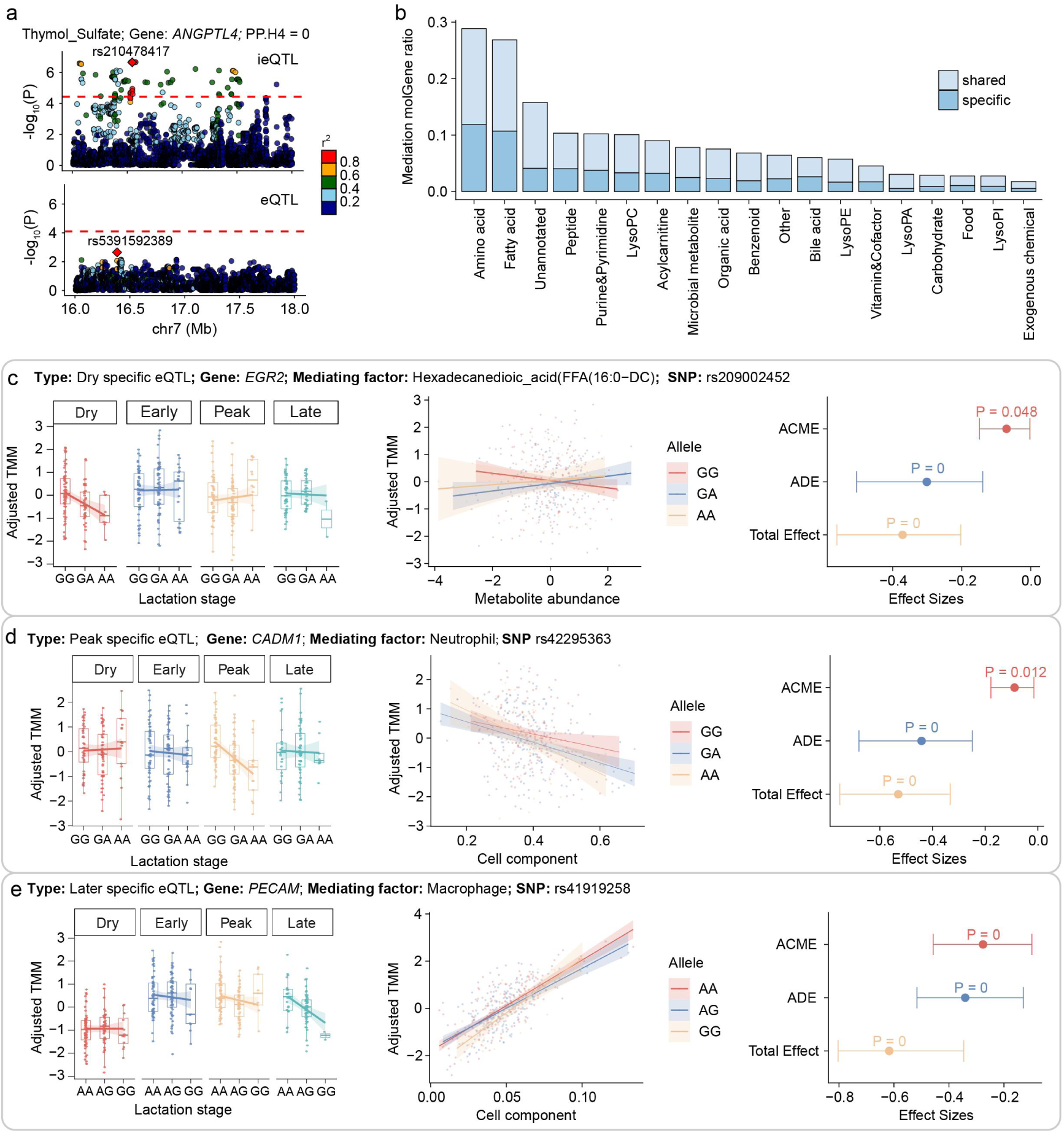
Mediation of lactation stage-specific molQTL by metabolite abundance and cell composition. **a**, Example of *ANGPTL4* regulated by Thymol sulfate interaction eQTL but not standard eQTL. **b**, GO enrichment analysis of differentially expressed genes (DEGs). **c-e**, Examples of stage-specific molGenes mediated by metabolite abundance and cell composition **(d,e)**.

**Extended Data Fig. 7.**
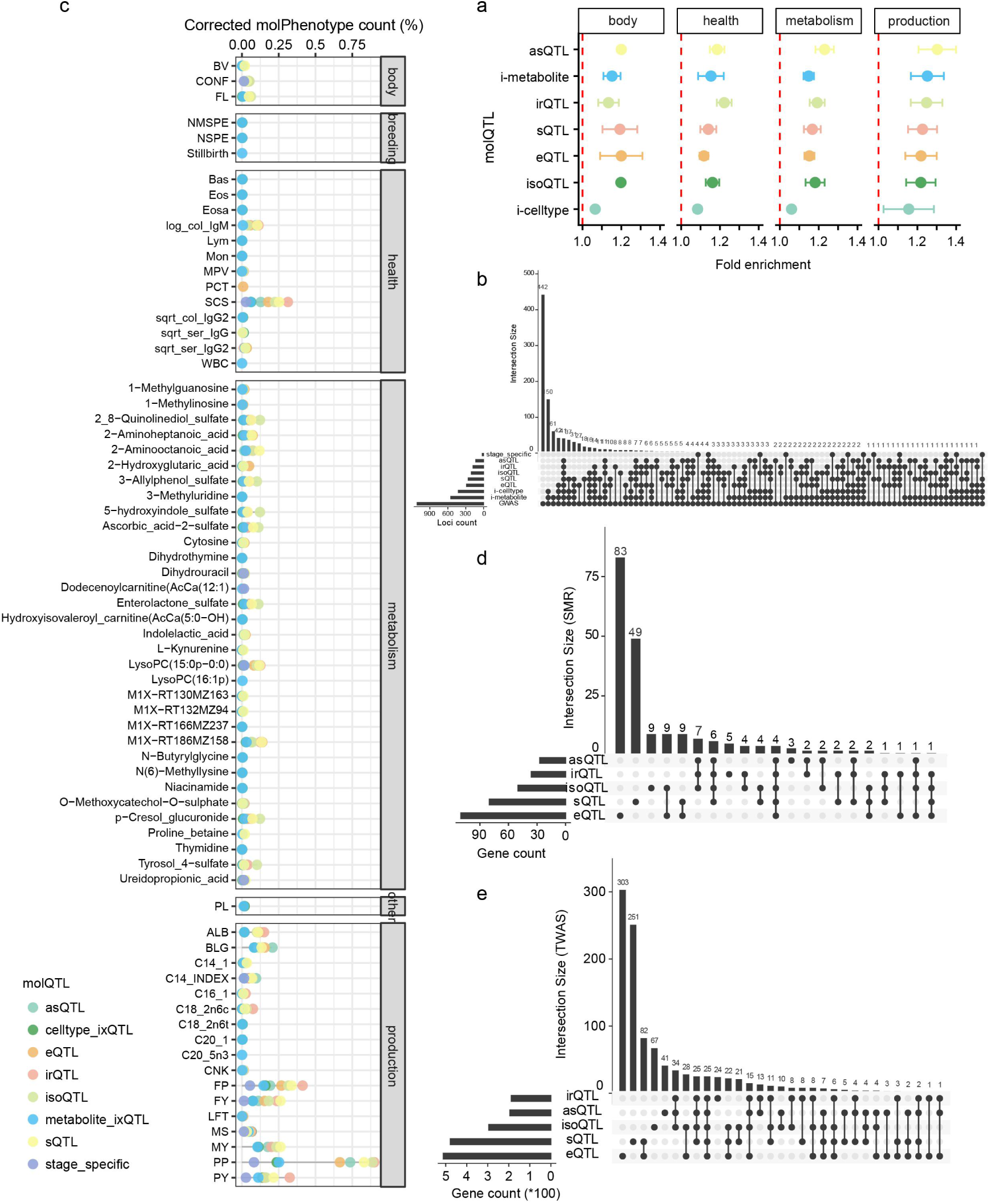
Complementarity of molQTL in interpreting GWAS loci. **a**, Enrichment (mean ± 95% confidence interval) of GWAS variants for five classes of standard molQTL and two interaction molQTL across four trait categories, including body conformation, health, metabolic and production. **b,** UpSet plot showing the number of GWAS loci colocalizing with at least one type of molQTL. **c,** Enrichment (mean ± 95% confidence interval) of GWAS variants for 11 types of molQTL. **d-e,** UpSet plot showing the number of trait-gene pairs identified using SMR **(d)** and TWAS **(e)** across molecular phenotypes.

**Extended Data Fig. 8.**
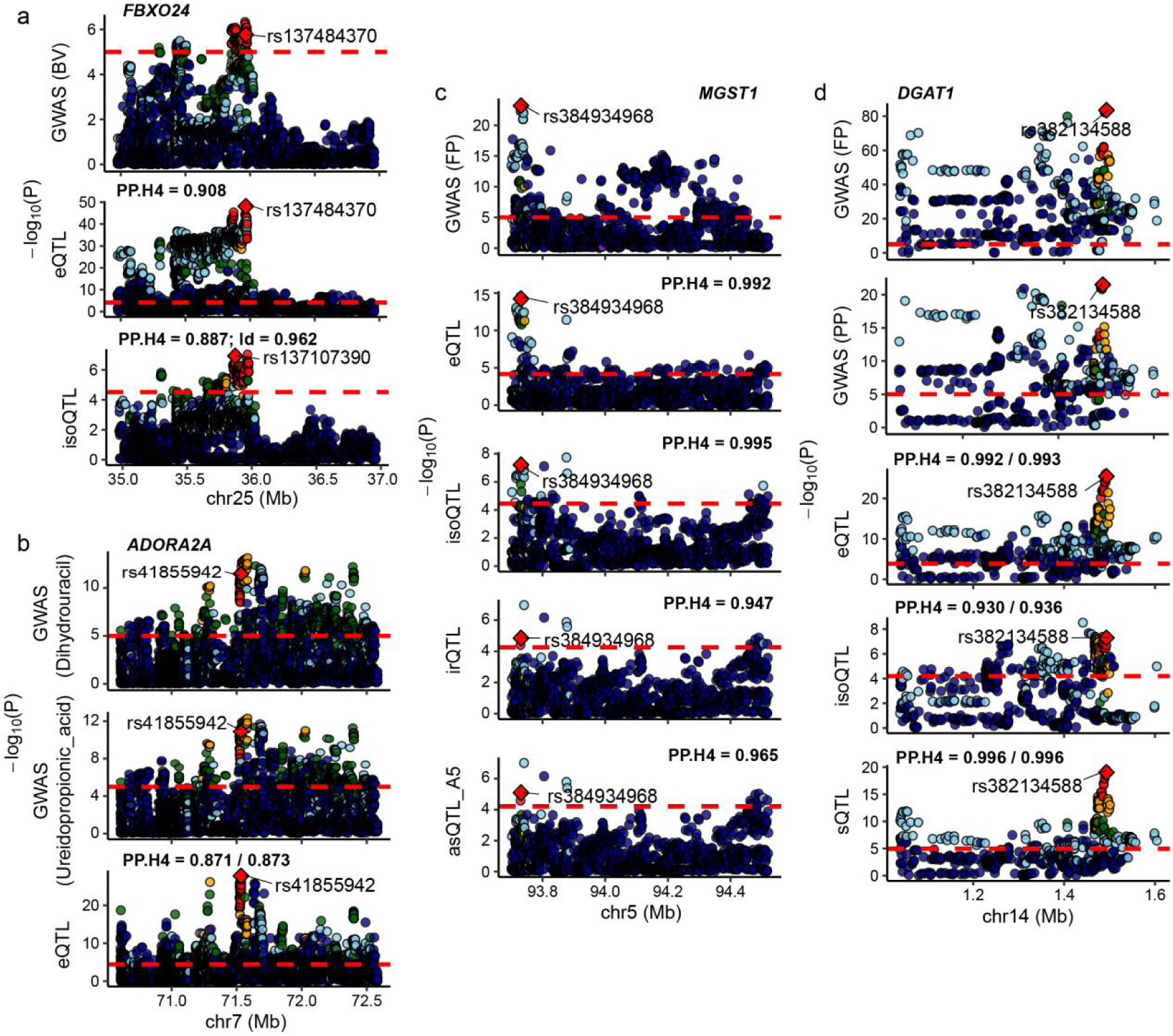
Examples of pleiotropic molQTL at complex trait GWAS loci. **a-d**, Colocalization results at four representative loci, including *FBXO24* (**a)**, *ADORA2A* (**b)**, *MGST1* (**c)** and *DGAT1* (**d)**), illustrating cases in which one or multiple molQTL regulate one or multiple complex traits.

**Extended Data Fig. 9.**
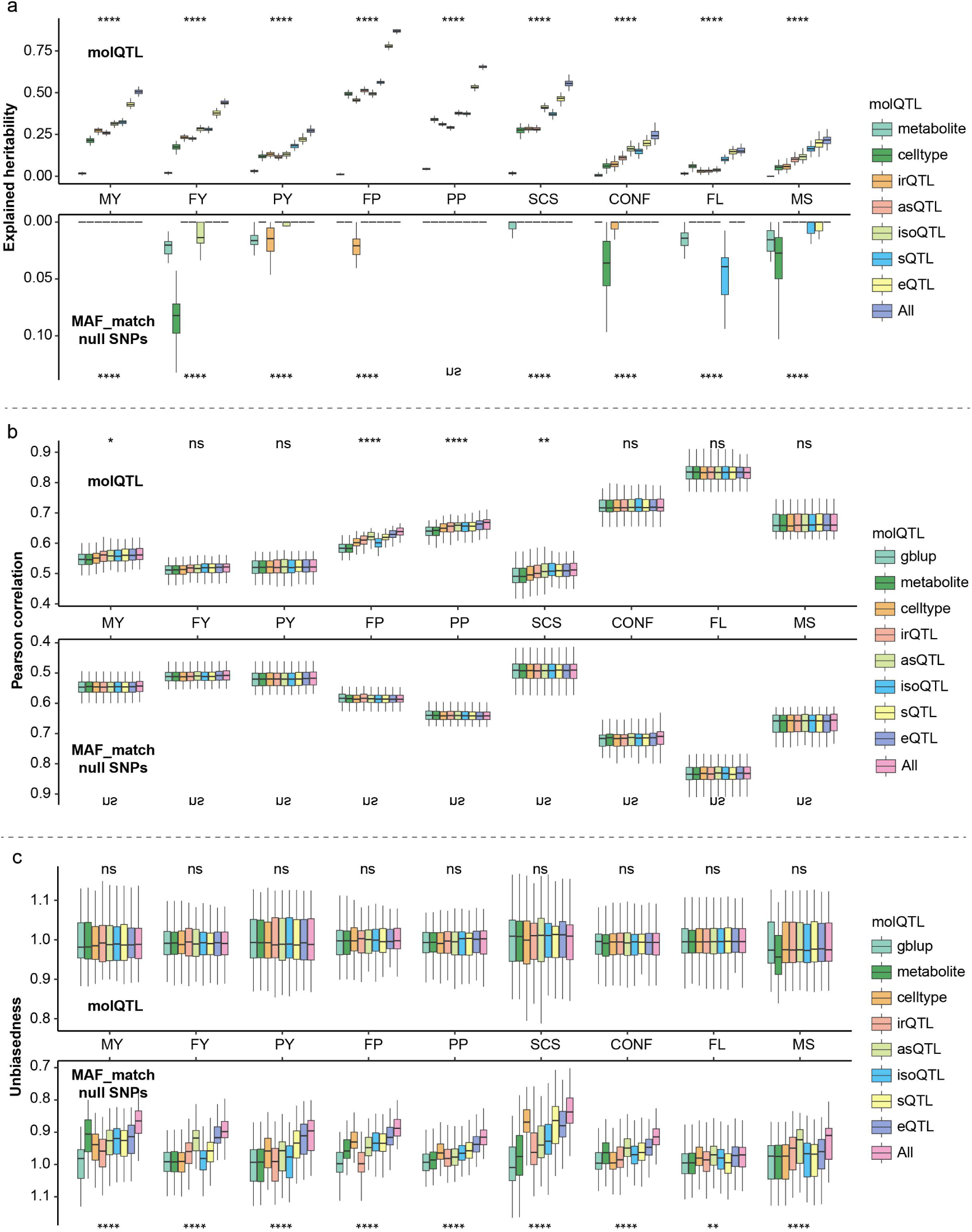
MolQTL improve genomic prediction of complex traits in dairy cattle. **a-c**, Box plots showing the contributions of seven classes of molQTL to nine major economic traits in dairy cattle, estimated using the GFBLUP model: proportion of heritability explained **(a),** genomic prediction accuracy **(b)**, and prediction unbiasedness **(c)**. molQTL-based models are shown at the top, and MAF-matched random variant models at the bottom. Box plots indicate the interquartile range (IQR) with the median, and whiskers extend to 1.5× IQR. P values were calculated using two-sided paired Wilcoxon signed-rank tests. ns, not significant; *, *P* < 0.05; **, *P* < 0.01; ***, *P* < 0.001.

## Notes

https://cattleblr.farmgtex.org

### Competing Interest Statement

The authors have declared no competing interest.

